# Scalable querying of human cell atlases via a foundational model reveals commonalities across fibrosis-associated macrophages

**DOI:** 10.1101/2023.07.18.549537

**Authors:** Graham Heimberg, Tony Kuo, Daryle DePianto, Tobias Heigl, Nathaniel Diamant, Omar Salem, Gabriele Scalia, Tommaso Biancalani, Shannon Turley, Jason Rock, Héctor Corrada Bravo, Josh Kaminker, Jason A. Vander Heiden, Aviv Regev

## Abstract

Single-cell RNA-seq (scRNA-seq) studies have profiled over 100 million human cells across diseases, developmental stages, and perturbations to date. A singular view of this vast and growing expression landscape could help reveal novel associations between cell states and diseases, discover cell states in unexpected tissue contexts, and relate *in vivo* cells to *in vitro* models. However, these require a common, scalable representation of cell profiles from across the body, a general measure of their similarity, and an efficient way to query these data. Here, we present SCimilarity, a metric learning framework to learn and search a unified and interpretable representation that annotates cell types and instantaneously queries for a cell state across tens of millions of profiles. We demonstrate SCimilarity on a 22.7 million cell corpus assembled across 399 published scRNA-seq studies, showing accurate integration, annotation and querying. We experimentally validated SCimilarity by querying across tissues for a macrophage subset originally identified in interstitial lung disease, and showing that cells with similar profiles are found in other fibrotic diseases, tissues, and a 3D hydrogel system, which we then repurposed to yield this cell state *in vitro*. SCimilarity serves as a foundational model for single cell gene expression data and enables researchers to query for similar cellular states across the entire human body, providing a powerful tool for generating novel biological insights from the growing Human Cell Atlas.

## INTRODUCTION

Characterizing the contexts in which cells employ different expression programs is critical for deciphering their functional role in health and disease. To date, well over 100 million individual cells have been profiled using single-cell or single-nucleus RNA-seq (sc/snRNA-seq) across homeostatic, disease, and perturbed conditions^1^. Individually, even the largest multi-tissue scRNA-seq atlases^2–4^ capture only a relatively small portion of cell states across human tissues; collectively, these atlases provide a vast, pan-human view of cell and disease biology that has the potential to address fundamental questions about human biology^1^. By aggregating across atlases, we may uncover biological insights and enable investigations into cell states that are common across multiple studies of the same organ and conditions (*e.g.*, similar neural progenitor populations across independent studies of brain development), different organs and conditions (*e.g.*, inflammatory fibroblasts in both ulcerative colitis and cancer); or between the human body and *in vitro* lab models (*e.g.*, regulatory T cells genetically perturbed to recapitulate *in vivo* cells from diseased tissue).

Despite this promise and the rapid growth in data, our ability to realize the potential of cross-datasets, pan-body, analyses remains limited and hampered by the need for laborious manual curation, harmonization, and dataset aggregation by expert analysts, as well as the painstaking process of selecting datasets, standardizing cell type annotations, and finding a common low-dimensional representation. As a result, most aggregation efforts have been limited in their biological scope and number of datasets, with some recent notable exceptions focused on genes rather than cell representation^5–8^.

To leverage and query the massive scale and richness of available single-cell atlases, we need both (1) a foundational model of cell states with an effective representation for single-cell profiles across different cell types and conditions that can be used across many applications without retraining; and (2) a measure of cell similarity that is robust to technical noise, scales to hundreds of millions of cells, and accurately generalizes to datasets and cell states not observed in the training. Established unsupervised methods to learn low dimensional representations of scRNA-seq profiles, such as Principal Component Analysis (PCA) or autoencoders^3,9–11^, faithfully preserve information from the input^3,9–11^ and may even eliminate technical variation for explicitly defined batches. However, they do not learn general features that encode relationships between cells needed to represent and query new data sets in the context of cross-study, pan-tissue biological variation.

Machine learning methods, metric learning in particular, have successfully learned representations for diverse entities and a measure of similarity between them, especially in image analysis. For example, metric learning models for facial recognition are explicitly trained to embed images of the same person closer together than images of different people, by exploiting visual features that are critical to distinguish individuals^12^. Once trained, images are embedded into a low-dimensional space, where distances between images represent a measure of similarity based on the learned features. Users can then query with an image not in the training set to find additional similar images that are nearby in the latent space and depict the same person. We reasoned that, analogously, metric learning could provide a meaningful representation of and similarity metric for cell profiles. By training a model using annotated scRNA-seq data, we can learn a low dimensional representation that places similar cells near each other and dissimilar cells farther apart. If learned from a sufficient diversity of cell profiles, such a representation would, in turn, provide a foundational model of cells and would allow efficient searches for cells with similar expression states (**Fig. 1a**).

**Fig. 1.**
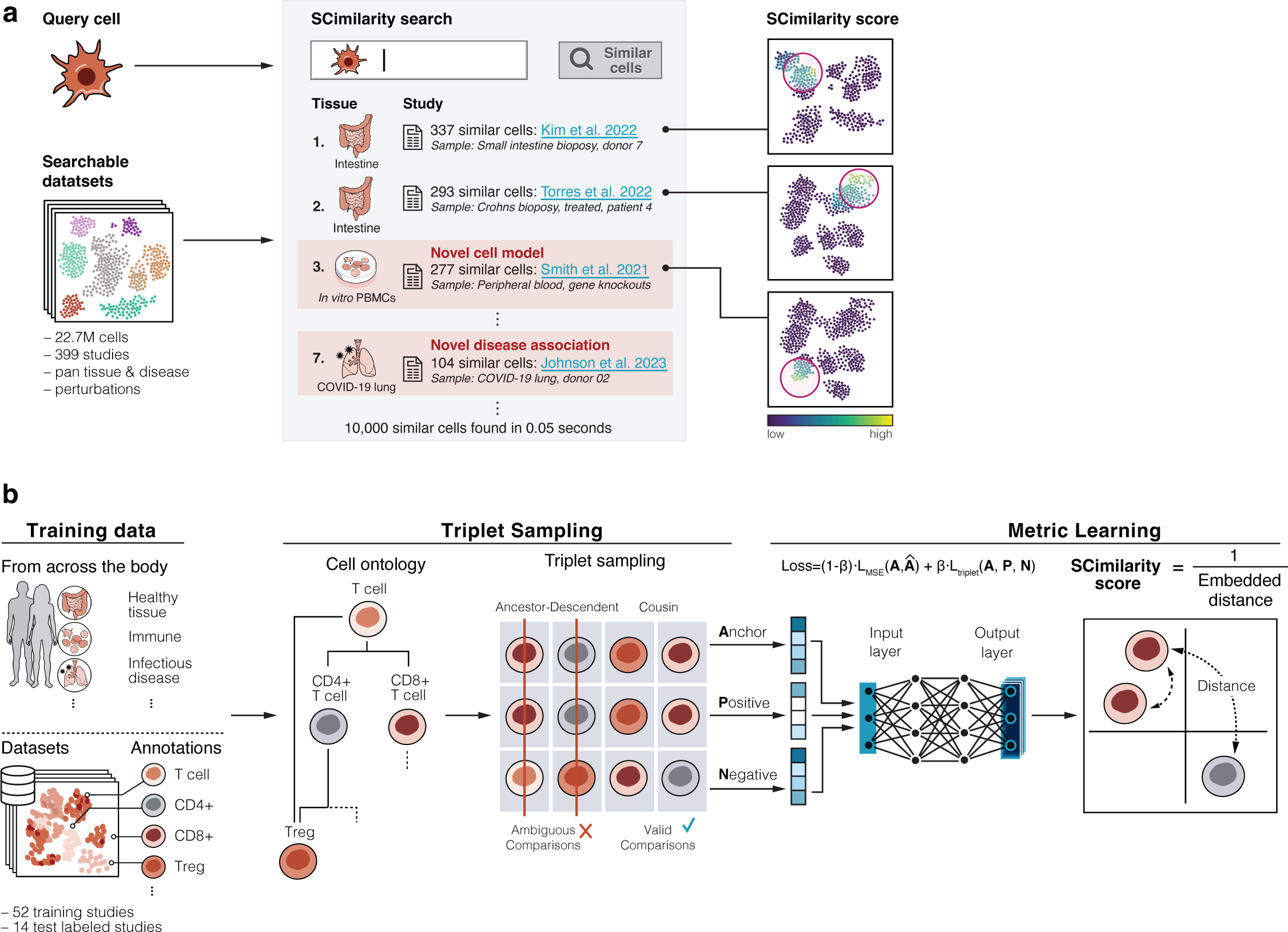
SCimilarity metric learning enables cell search in large human scale atlases. **a,** Cell Querying with SCimilarity. Left: A query cell profile is compared to a searchable reference collection of 22.7M profiles from 399 studies. Center: Sample with similar cells are identified and returned with information about the original sample conditions, including unexpected tissue, *in vitro* or diseases contexts. Right: A SCimilarity score is computed between the query cell and each cell within a tissue sample. **b,** Triplet loss training. From left: 52 training and 14 test annotated (by the Cell Ontology) datasets from across the body are sampled for cell triplets (an anchor, a “positive” (anchor-similar), and a “negative” (anchor-dissimilar) cell; based on Cell Ontology annotations) to train a neural network that embeds similar cells closer than dissimilar ones (**Methods**).

Here, we introduce SCimilarity, a class of deep metric learning models that quantify similarity between single-cell expression profiles (SCimilarity score) and provide a single-cell gene expression foundational reference model to systematically query for comparable cell states across tissues and diseases. SCimilarity uses a training set of diverse author-annotated cell profiles to learn a universal representation and distance metric that facilitates efficient searches across a massive reference meta-atlas for the most expression-similar cells. To train a foundational model that can be broadly applied to many applications across tissues and studies, we built a programmatic pipeline for massive data import and automated standardized curation and used it to assemble a corpus of 22,699,774 cells from 399 datasets spanning a broad range of organs, systems and conditions across the human body. After training and testing on a subset of 66 studies and 7.9M single-cell profiles, the learned models generalize well, representing and quantifying similarities between 14.9M cells from another 347 studies excluded from training. By tuning a single parameter during SCimilarity’s training, we yield models optimized for either data integration and visualization of millions of cells across hundreds of studies, or for fast and efficient (millisecond) queries of a new cell state across tens of millions of cells. Finally, we illustrate the power of SCimilarity by querying for a fibrosis-associated macrophage (FMΦ) subset previously identified in interstitial lung disease (ILD), finding comparable cell populations (but with different annotated names and signatures) in other ILD studies, as well as in new contexts, including COVID-19, different tumors including pancreatic ductal adenocarcinoma (PDAC), and even healthy lung (at low abundance). Surprisingly, SCimilarity recovered FMΦ-like cells among PMBCs cultured and stimulated in a 3D hydrogel system *in vitro*, which we experimentally validated, producing an FMΦ-like *in vitro* cell system for future functional studies of a tissue-resident cell state. Overall, SCimilarity preserves expression diversity across cells in an integrated foundational model of a human cell atlas, and allows a novel scaled cell search across organs, systems, and conditions, as a powerful framework for generating biological insights and experimentally testable hypotheses.

## RESULTS

### SCimilarity: novel similarity metrics and representations for single-cell expression profiles

SCimilarity is a family of models that blend unsupervised representation learning and supervised metric learning, through simultaneously optimizing two objectives : (1) a supervised triplet loss function, which is used to embed expression profiles from matching cell types close together, effectively integrating cells of the same type across studies^13–15^, and (2) an unsupervised mean squared error (MSE) reconstruction loss function, which encourages the model to preserve variation from the input expression profiles, capturing subtler differences in expression patterns within cells of the same type, such as those related to tissue residency of immune cells (**Fig. 1b**, **Methods**). The balance of these two objectives, set by a single hyperparameter, β, determines the properties of the representation (**Methods**). Increasing the relative weight of the triplet loss function improves dataset integration, while increasing the relative weight of the reconstruction loss improves querying performance. Therefore, different loss function weightings within the same model architecture can address different applications.

We train SCimilarity with tens of millions of cell triplets sampled from data with author-provided standardized cell type annotations from the Cell Ontology^16^ (**Fig. 1b**, **Methods**). Specifically, each training triplet consists of similar anchor and positive cells (*i.e.*, same cell type) from different studies, while anchor and negative cells are dissimilar (*i.e.*, distinct cell types; from the same or a different study). However, even with standardized Cell Ontology terms, some cell type comparisons are ambiguous due to arbitrary differences in annotation granularity across studies (*e.g.*, it is ambiguous if cells annotated as “T cell” in one study and “CD4^+^ T cell” in another are similar or dissimilar). To address this, SCimilarity excludes cell pairings with such vertical ancestor-descendant Cell Ontology relationships from training triplets, and learns only from cells that are either explicitly similar or unambiguously dissimilar (**Fig. 1b**, **Methods**). By sampling only unambiguous triplets we eliminate the need to manually flatten or harmonize every cell type annotation and are able to seamlessly scale the training set across dozens of studies.

### A learned SCimilarity representation of 22.7M cells across dozens of tissues and disease datasets collated by an automated curation and processing pipeline

To test SCimilarity models, we assembled a compendium of sc/snRNA-seq datasets across human biology. We focused on studies generated with one experimental platform (10x Genomics Chromium droplet-based scRNA-seq) and data publicly available on the Gene Expression Omnibus (GEO)^17^ or CELLxGENE^18^. These data capture much of the published scRNA-seq data, and were generated with similar library preparation protocols and computational pipelines^19^. There were 753 human sc/snRNA-seq datasets matching our search criteria and keywords as of March 23^rd^, 2021 (with Biopython Entrez^20^, **Methods**). The number of samples and cells matching our criteria has at least doubled every 6 months between December 2018 and March 2021; (**Extended Data Fig. 1a,b**). We programmatically downloaded 13,401,599 cell profiles from 333 of the identified studies with their respective GEO metadata and unnormalized gene count matrices (**Methods, Extended Data Table 1**). We manually ingested another 66 well-annotated studies from either the CELLxGENE portal^18^ or from large studies and consortia not available through GEO that passed the same dataset filtering criteria (**Methods**). Overall, we assembled a corpus of 399 studies comprising 22,699,774 cells from 33,815 tissue samples with 184 unique Tissue Ontology terms^21^, 132 Disease Ontology terms^22^, and 204 Cell Ontology cell type terms^16^, with each Cell Ontology term appearing in at least two separate datasets (**Fig. 2a**, **Extended Data Fig. 1c**, **Extended Data Table 1**).

**Fig. 2.**
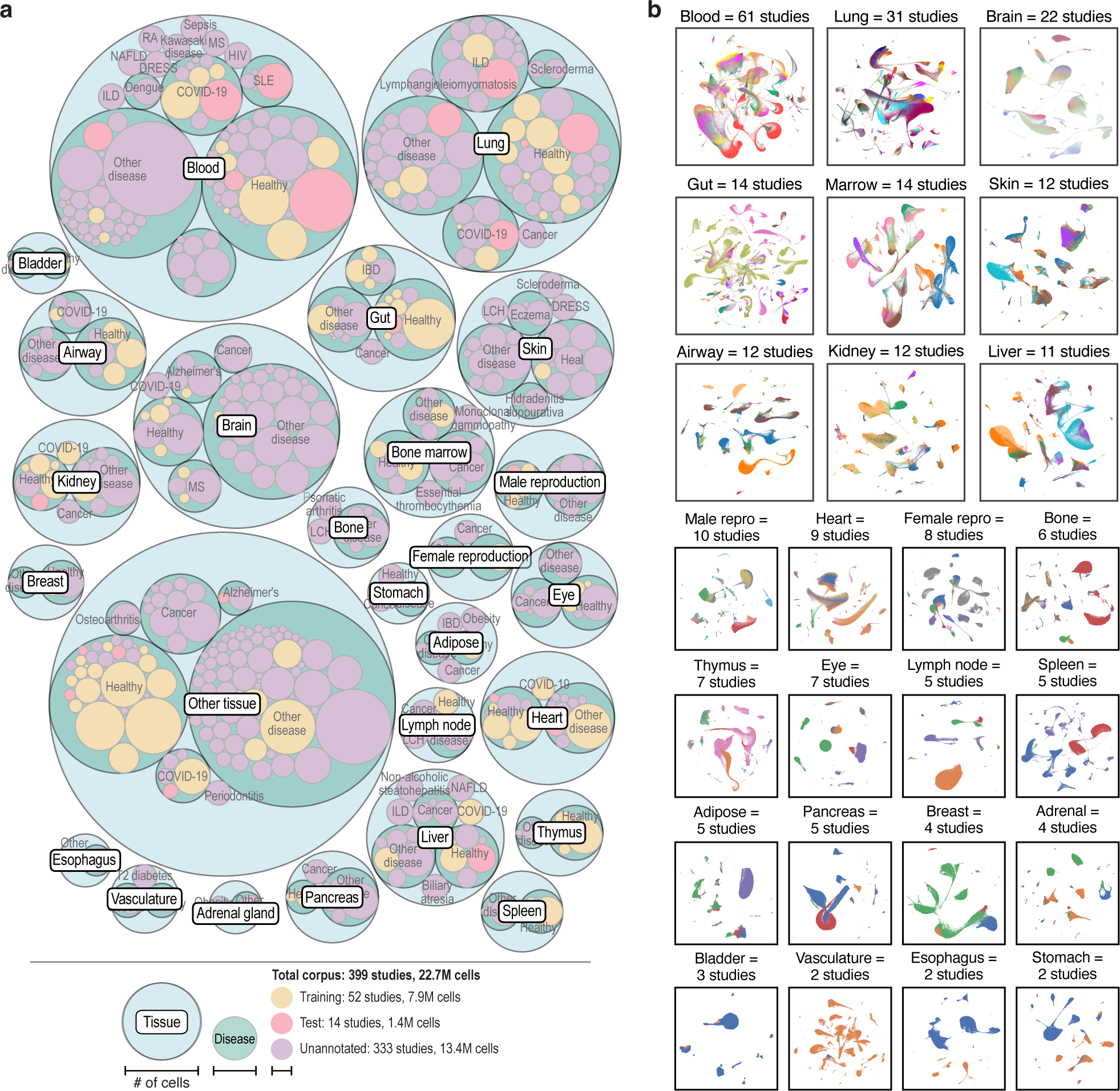
SCimilarity learns a universal representation that generalizes to new datasets. **a,** A large-scale reference database of public gene expression datasets across tissues and diseases. Number of cells (circle size) across tissues (outermost light blue circles) and disease states (middle green circles) across individual studies (innermost circles) in the training (gold), test (pink) or unannotated (purple) datasets. **b,** Integration between studies without feature selection or batch correction. Uniform manifold approximation and projection (UMAP) embedding of cell profiles (dots) generated on a 128-dimensional latent space from SCimilarity’s integration model (**Methods**) for cells from 21 tissues (panels) and 239 unique studies (color code). For tissues with more than a million cell profiles, the UMAP embedding was computed on a random uniform subsampling of 1 million cells from the studies for that tissue.

We trained SCimilarity models with a training set of 7,913,892 single-cell profiles from 52 studies with Cell Ontology author annotations that reflected a diversity of conditions and tissues (**Extended Data Fig. 1d**, **Extended Data Table 1**), sampling 50,000,000 of the most informative triplets (**Methods**). We withheld 14 studies comprising 1,384,283 cells with Cell Ontology annotations for testing the learned representation and metric (**Fig. 2a**). We excluded tumor, cell lines, and iPSC-derived samples from the training and test sets, because cell identity of tumor cells and cell lines can be ambiguous.

### Tuning of SCimilarity’s reconstruction and triplet loss functions yields models optimized for integration *vs*. cell search tasks

We examined six different blends for SCimilarity’s objective function, varying the relative weighting of the reconstruction and triplet loss functions, and finding that the two loss function components gave rise to different behaviors in a trained model. Briefly, we assessed the models on two tasks – data integration and searching for cells similar to a query profile – using studies entirely held out from training. To evaluate data integration, we quantified how coherently cells of each type are clustered and how distinct each cell type cluster is from other clusters. To this end, we created an ontology-aware variation of average silhouette width^23^ to quantify integration capabilities across datasets without harmonizing cell type annotations (**Methods**). To evaluate our cell search distance metric, we compared searches with SCimilarity to gene signature scoring (**Methods**). The higher the correlation between these two quantities, the more our similarity metric corresponds to traditional signature-based similarity to represent a cell state of interest.

Models with higher triplet loss weighting scored higher on integration benchmarks, while models with higher reconstruction loss weighting encoded distances between cells in a manner better correlated with differences in representative expression signature scores (**Extended Data Fig. 2a,b**). Pure triplet loss, which is calculated at the level of cell type labels, does not reliably preserve subtle cell state differences, such as tissue specificity or disease response within cells of the same type. Mean squared error reconstruction loss complements this by preserving more subtle gene expression patterns, while the triplet loss ensures that cells of the same type are embedded closely together. Based on the biological question, a user can tune this balance to yield the highest utility. We thus pursued two SCimilarity models: an integration model, optimized for the task of learning a low dimensional representation that groups cells by type rather than by study; and a cell search model that is optimized for the task of retrieving cells with an expression state similar to that of a query cell across hundreds or thousands of scRNA-seq datasets (**Extended Data Fig. 2a**, **Methods**).

### SCimilarity’s latent space representation filters outlier cells and integrates test datasets without batch correction

We next benchmarked if SCimilarity’s latent space representation from the integration model generalizes well to cells from entire datasets held out of training compared to other methods. In a low-dimensional embedding, unannotated cell profiles from nine lung studies (7 training set, 2 test set) visually intermix well when embedded into SCimilarity’s learned 128-dimensional space (**Extended Data Fig. 2a**). SCimilarity’s data integration model scored higher than Harmony, scVI, and scArches on integration tasks by the ontology-adjusted ASW measure of cluster coherence, but scored lower for normalized mutual information (NMI) and adjusted rand index (ARI), which measure the extent of study mixing within each cluster (**Extended Data Fig. 2b**). Thus, without directly training on the full data set or performing additional batch correction, the integration model clusters cells by type rather than study at a level that is competitive with existing methods trained directly on the data. This demonstrates that the triplet loss learns features that capture meaningful biology, while reducing technical sources of noise and avoiding overfitting to the training set.

SCimilarity quantifies a confidence level for each cell’s representation, providing both outlier detection and an assessment of the representation’s relevance in the context of new data. When computing the representation of a new cell, the further outside the scope of model training it is, the harder it is for the model to accurately represent it. Using SCimilarity’s score to quantify how distant a query cell is from the training data distribution provides a heuristic about the quality and scope of the representation – a cell scoring as highly similar to cells seen during training can be confidently represented by the model. Overall, 79.5% of *in vivo* holdout cells had high representation confidence. Tissue samples with particularly low representation confidence, such as stomach (n = 0 training studies), fetal gut (n=1), and bladder (n=0) were either absent or poorly represented in training (**Methods, Extended Data Fig. 2c**), suggesting that more labeled training datasets from those tissues could improve the model’s representation. Similarly, 43.8% of *in vitro* cell profiles were considered low confidence due to poor matching to the training set (which excluded *in vitro* samples).

We combined SCimilarity’s ability to generalize to new datasets and its confidence-based filtering to systematically generate meta-atlases for 21 different human tissues without labor-intensive dataset harmonization and no additional training (**Fig. 2b**). If datasets have already been embedded using SCimilarity, this task only requires concatenation of cells of interest and standard visualization workflows.

### SCimilarity assigns an unannotated query cell to a cell type by finding similar cells in a labeled reference

We next used SCimilarity to find the cells in the annotated reference that are most similar to an unannotated query cell profile, and then annotate the query cell accordingly (**Fig. 3a**, **Methods**). This approach is distinct from established annotation methods in that it (1) relies on a large, pan-human annotated cell repository, (2) employs a measure of expression similarity, and (3) classifies at the single cell rather than cluster level, providing greater transparency into the classification itself. Thus, users can see which individual cells, studies, and tissues are driving the classification decision. Moreover, since each cell is annotated independently, no clustering or associated parameter selection, such as the number and resolution of clusters, are required. A user can choose to annotate a cell’s profile by comparing it either to a desired subset of cell types (*e.g.*, for a tissue-specific query) or to the entire annotated cell reference. Because SCimilarity is built using metric learning, finding the most similar cells is the same as retrieving the query cell’s nearest neighbors. This operation is extremely efficient with the hnswlib algorithm^24^, where searching a precomputed approximate nearest neighbor index of all the annotated reference cells in SCimilarity’s latent space takes just 20 milliseconds (**Methods**). Low SCimilarity scores to reference cells flag an outlier query cell, which may be either a cell type that is not within the reference or a query cell of low quality.

**Fig. 3.**
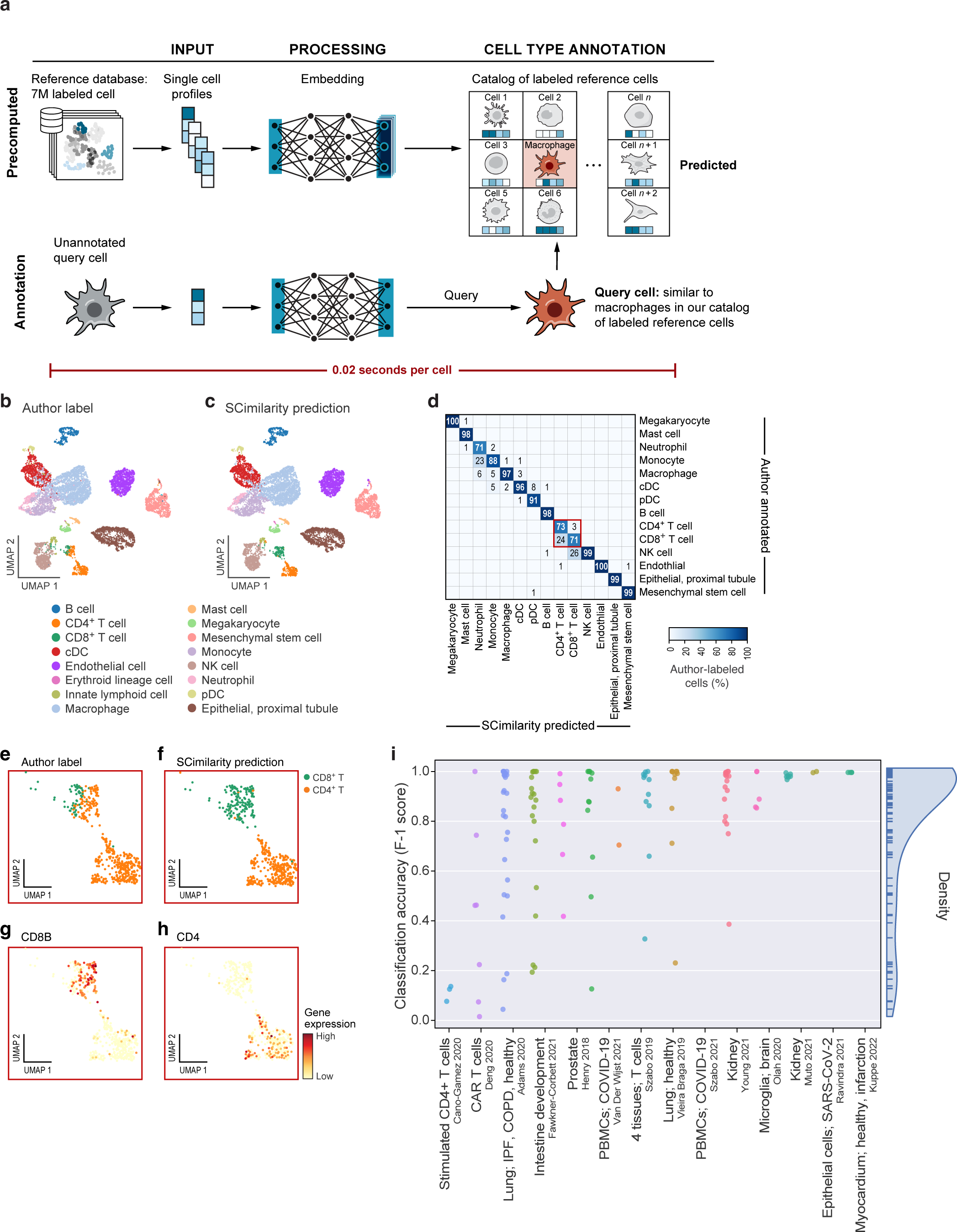
SCimilarity accurately annotates cell types across the human body. **a**, SCimilarity cell annotation. A new unannotated cell (red, bottom left) is embedded in SCimilarity’s common low-dimensional space and compared against the precomputed reference for cell type annotation (0.02 seconds per cell). **b-d**, SCimilarity annotation of a kidney scRNA-seq dataset. (**b,c**) UMAP embedding of cell profiles (dots) from SCimilarity’s latent representation of a held out kidney dataset^1^ colored by author provided (**b**) or SCimilarity predicted (**c**) cell type annotations. (**d**) Percentage (color bar and number) of author-annotated cells (columns) with each SCimilarity annotation (rows). **e-h,** SCimilarity-corrected author annotations. UMAP embedding of cell profiles (dots) from SCimilarity’s latent representation for cells either author-annotated or predicted as CD4^+^ or CD8^+^ T cells, colored by author (**e**) or SCimilarity (**f**) annotations, and by *CD8A* (**g**) or *CD4* (**h**) expression. **i,** Classification performance. F1 score (y axis) for SCimilarity *vs*. author annotation for each cell population (dot) from each of 14 held out datasets (x axis). Right: Distribution of F1 scores.

SCimilarity quickly and accurately assigned cell types for entire datasets held out from training, as well as for the rest of the 22.7M cell corpus. When limiting potential cell types to author-selected labels, 94.5% of SCimilarity’s predicted labels from healthy kidney samples^25^ match the author-provided cell type annotations (**Fig. 3b-d**, **Methods**). In some cases, where SCimilarity’s predictions did not match author-provided annotations, SCimilarity’s predictions were more accurate or granular. For example, 94% of the cells that the authors^25^ annotated as CD4^+^ T cells but SCimilarity annotated as CD8^+^ T cells express *CD8A* or *CD8B* (and none express *CD4*), supporting SCimilarity’s annotation (**Fig. 3e-h**). Separately, when allowing cells to be annotated as any cell type in the repository, 6.3% of the author-annotated CD4^+^ T cells were reannotated by SCimilarity as regulatory T cells (T_regs_) (**Extended Data Fig. 3a**), most of which (85.2%) expressed at least one T_regs_ marker (FOXP3, IL2RA, or IKZF2, **Extended Data Fig. 3b-d**). Similarly, 1.8% of author-annotated mesenchymal stem cells (**Fig. 3b**) were reassigned by SCimilarity as myofibroblasts (**Extended Data Fig. 3a**) and 93% of those express the myofibroblast-associated gene *ACTA2* (**Extended Data Fig. 3e**). Cell type prediction was rapid, taking 3-5 seconds to embed and annotate 10,000 cells from a dataset. Overall, across all 14 test datasets spanning 78 Cell Ontology terms, 71% of the cell populations had high agreement (>85% of the cell population) between author and SCimilarity annotations (**Fig. 3i**). SCimilarity performed poorly on one dataset (Cano-Gomez et al.^26^), due to fine granularity and redundancy of author labels (*e.g.*, CD4^+^ ɑβ T cells, helper T cell, memory T cell, naive T cell, and regulatory T cell).

We used SCimilarity’s cell type assignment to rapidly annotate all 22.7M cell profiles in one common model, newly-annotating 13,401,599 profiles and reannotating 9,298,175 author-annotated profiles (**Methods**) to a single set spanning 74 cell type labels (21 coarser lineages) from 25 simplified tissue categories (**Fig. 4a,b**, **Extended Data Fig. 3f**). A consistent annotation across datasets facilitates cross-study and cross-tissue analyses of one cell type or lineage, as SCimilarity can extract cells from hundreds of studies, aggregating vast biological diversity across one cell type. For example, we readily aggregated 1,172,325 fibroblasts and myofibroblasts (**Extended Data Fig. 3g**) and 2,507,879 monocytes and macrophages (**Extended Data Fig. 3h**) from hundreds of studies profiling different primary tissue samples.

**Fig. 4.**
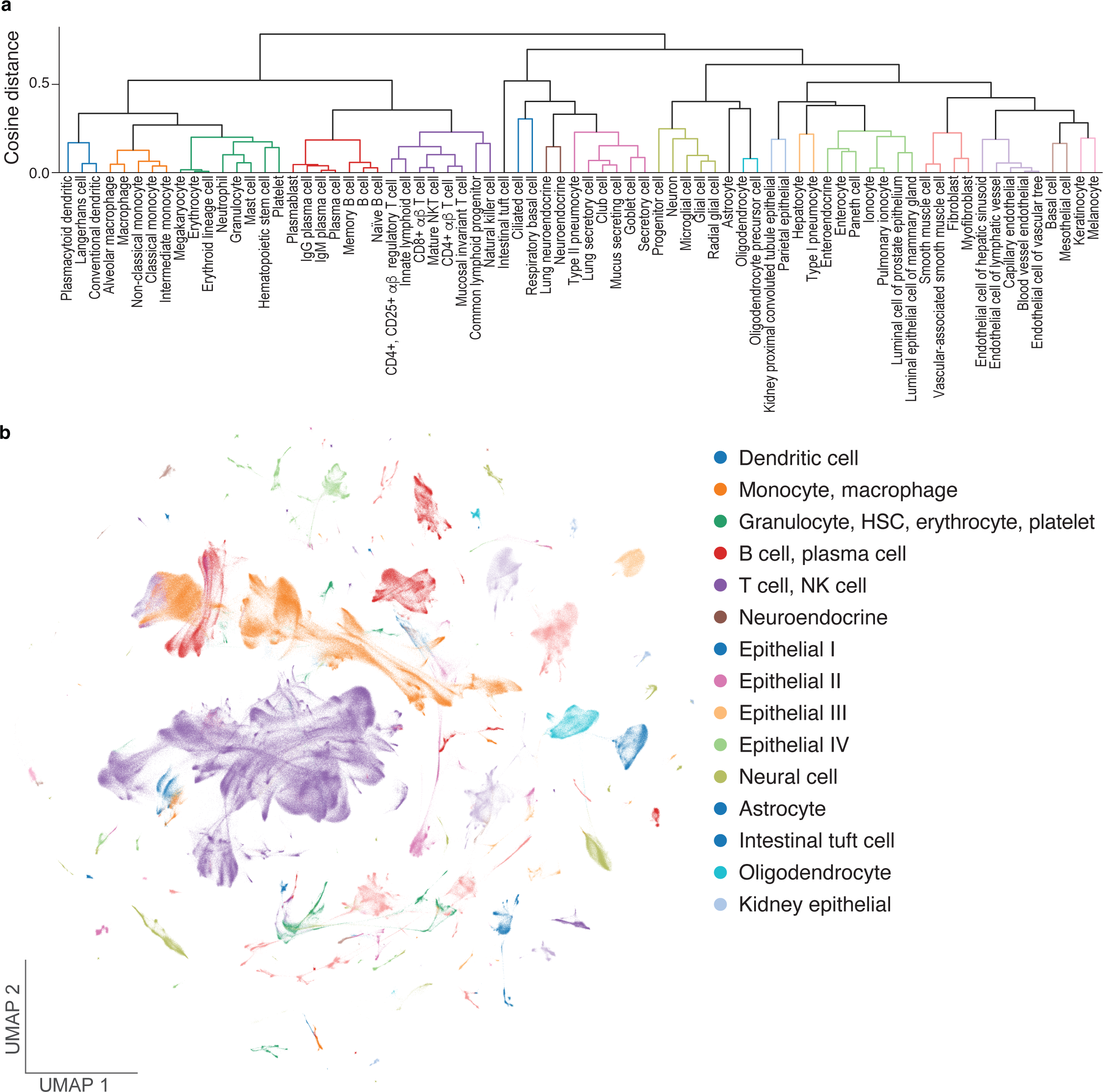
SCimilarity annotations scale to tens of millions of cells from hundreds of datasets. **a,b** Predicted cell types group by biologically-accurate lineages. **a,** Hierarchical clustering dendrogram of centroids of predicted cell types (leaves) in SCimilarity latent space, colored by lineage. Clustering was performed using cosine distance and average linkage. **b,** UMAP of 2,000,000 embedded cells uniformly sampled from the 22.7M reference, colored by clusters (as labeled in **a**).

### SCimilarity’s representations comprise of interpretable biological features

To interpret SCimilarity’s annotations, we quantified the importance of each gene for cell type annotations assigned by the foundational query model using Integrated Gradients, a method that identifies the impact on model predictions of small disturbances to the input expression profiles (**Methods**). For example, the top gene attributions that distinguish lung alveolar type 2 (AT2) cells are surfactant genes *SFTPA2*, *SFTPA1*, *SFTPB*, and *SFTPC*, consistent with known AT2 cell function^27^. SCimilarity learned these without prior knowledge of cell type specific genes, signatures, or highly variable genes. Overall, SCimilarity’s top importance genes agreed well with differentially expressed marker genes for 17 different matched types^3^ with the exception of rare neuroendocrine cells (average AUC=0.84, **Extended Data Table 2, Extended Data Fig. 3i**). Thus, SCimilarity’s representation captured known and validated biological markers within its features.

### Cell search identifies fibrosis-associated macrophages across tissues and diseases

With a single representation and common definition of cell types, we hypothesized that SCimilarity could help elucidate the role of tissue-resident immune cells. As a case study, we focused on macrophages, given their remarkable plasticity in cell states and their important specialized roles in tissue repair, regeneration, and fibrosis^28,29^. Recent scRNA-seq studies in fibrotic diseases, including lung fibrosis, cancer, obesity, and COVID-19 have reported seemingly-similar *SPP1*^+^ fibrosis-associated macrophage (FMΦ) populations^30–38^. However, because each study identified them independently, using different nomenclatures and marker gene signatures to define subsets, it is unclear how similar these cell states are. Moreover, it is unknown how broadly associated such cell states are with other diseases, especially those with prominent fibrosis. We reasoned that SCimilarity’s cell search should allow us to query our corpus with an FMΦ cell profile from one study to identify similar cells across other tissues and conditions, thereby clarifying the cell identity of similarly-described cells and the conditions in which FMΦ arise (**Fig. 5a**).

**Fig. 5.**
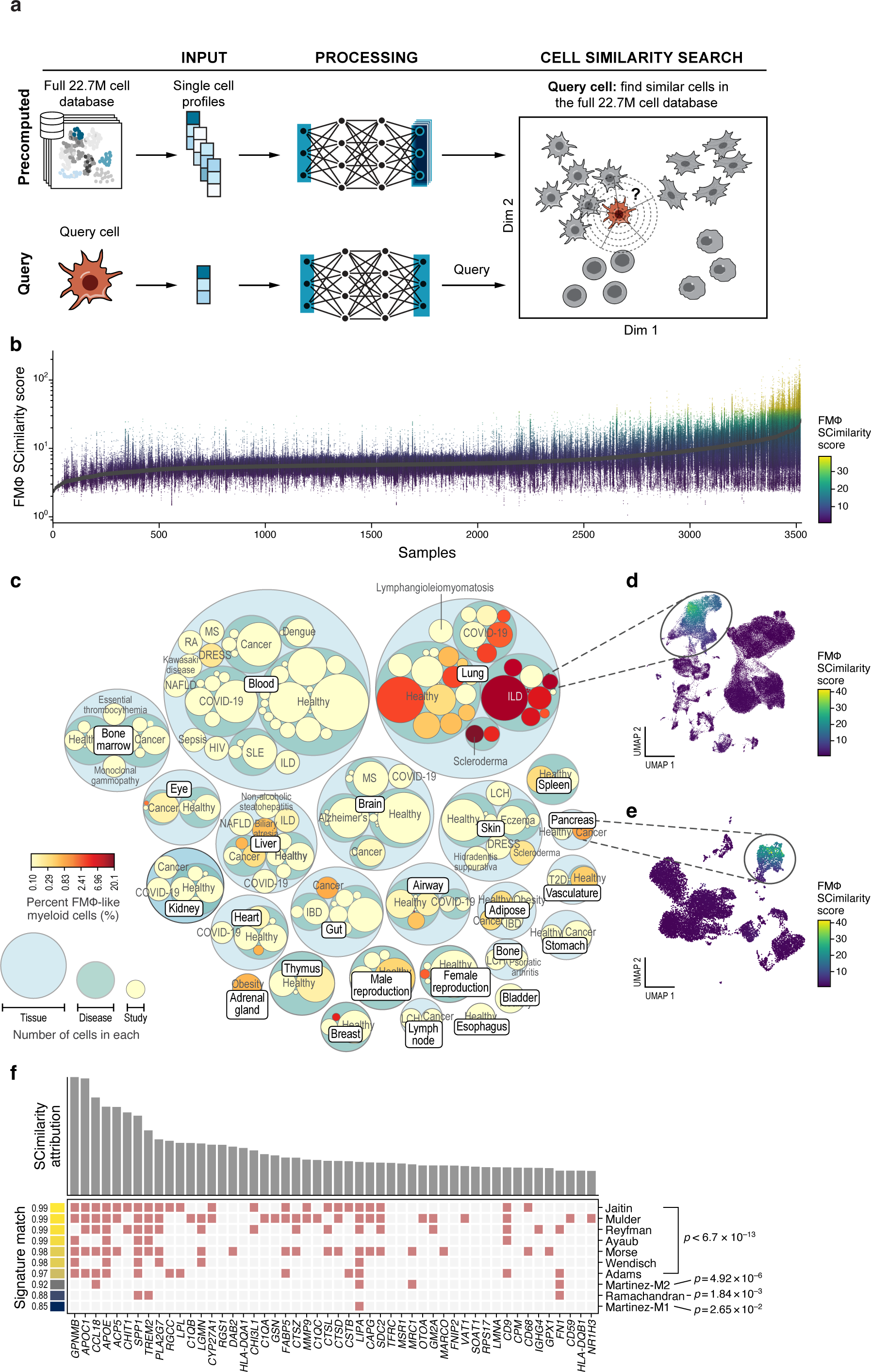
SCimilarity cell search reveals FMΦs across ILD and other diseases. **a,** SCimilarity cell search. A query cell profile (bottom left) is embedded into the learned SCimilarity representation along with the reference of 22.7M cells, and its nearest neighbors, determined by distance from the embedded query in the low dimensional space, are tabulated by study, tissue and disease. **b-e,** Identification of FMΦs across tissues by SCimilarity cell search. (**b**) SCimilarity scores (y axis, log_10_ scale, and color bar) against a FMΦ query profile for each annotated monocyte or macrophage (dot) from 1,041 *in vivo* tissue samples from 143 studies (x axis), ordered by mean SCimilarity score. (**c**) Number of cells (circle size) across tissues (outermost light blue circles), disease states (middle green circles), and individual studies (innermost circles, colored by fraction of monocytes and macrophages with SCimilarity scores >95^th^ percentile of all FMΦ SCimilarity scores (log scaled color bar)). Circle size for disease and individual study are scaled relative to other diseases in the same tissue, or studies in the same disease. (**d,e**) UMAP embeddings of cell profiles (dots) from the SCimilarity representation (query model) from an ILD^2^ (**d**) and PDAC^3^ (**e**) studies, colored by FMΦ query SCimilarity scores (color bar). **f,** Identification of FMΦs associated genes by importance. Integrated gradients attribution scores (y axis, top) for the genes (x axis top, and columns, bottom) with top 50 scores for FMΦs *vs*. lung macrophages (**Methods**), and their membership (red: presence; grey: absence) in published macrophage signatures (bottom, rows). Left color bar: AUC of the ranking of each published signature in SCimilarity attribution scores (AUC=1: all *n* signature genes are listed as the top *n* genes by SCimilarity attribution scores for distinguishing FMΦ). Martinez-M1 and -M2: macrophage states expected to be different from FMΦs. P-value: hypergeometric test

We queried our model with the FMΦ cell profile, searching for similar cells across 2,578,221 cells annotated by SCimilarity as monocytes or macrophages in the 22.7M cell corpus (**Fig. 5a**). SCimilarity queries can use either an individual cell profile or a centroid of multiple cell profiles. Here, we input the centroid profile of a macrophage cell subset from Adams *et al.*^30^ that we defined using a gene signature consisting of the extracellular matrix remodeling and fibrosis-associated genes *SPP1*, *TREM2*, *GPNMB*, *MMP9*, *CHIT1*, and *CHI3L1* (**Methods**). In two seconds, SCimilarity exhaustively computed the pairwise similarity of our query profile to each of the 2.6M *in vivo* profiles of the cells it annotated as monocytes or macrophages in our corpus (**Fig. 5b and Extended Data Fig. 4a**). Alternatively, simply identifying the 10,000 cells with the highest SCimilarity score takes 0.05 seconds (**Methods**). By comparison, a more conventional approach that scores each cell in the corpus with a literature-defined FMΦ gene signature took 2 hours and 46 minutes (**Extended Data Fig. 4b**). The gene signature and SCimilarity scores are broadly correlated (𝑟 = *0*.*50*, 𝑝 < *10*^−300^, **Extended Data Fig. 4a-c**), showing that the granular cell state, not just the cell type, is well-represented in SCimilarity query score and embedding.

The SCimilarity search showed that FMΦs are common in ILD lung samples in our compendium, as well as present in some cancers, including uveal melanoma, pancreatic ductal adenocarcinoma (PDAC), and colon cancer (**Fig. 5c-e**, **Extended Data Table 3**). Of the top 1% of monocytes and macrophages most similar to our query, 99.1% were from lung tissue and 87.2% from ILD and COVID-19 lung samples. The prevalence of FMΦ-like cells in the lung varied by disease: the proportion of monocytes and macrophages that were FMΦ-like was 20% and 4% in two systemic sclerosis (SSc) studies, 6.1% on average (SE = 1.4%) across 13 ILD studies (excluding SSc), 1.2% on average across seven COVID-19 lung studies (SE = 0.5%, 0% in non-lung COVID-19 data) and 0.4% in 19 studies annotated as “healthy”, “normal” or with no disease annotation (SE = 0.2%). While abundant in SSc lung, FMΦ-like cells were much rarer (0.14% of myeloid cells) in SSc skin^39^. There were some FMΦ-like cells in other fibrotic diseases and tissues, such as one primary pancreatic ductal adenocarcinoma (PDAC) tumor^40^ (0.85% of 1,171 myeloid cells) and one liver metastasis^41^ of PDAC (0.5% of 1,199 cells). Thus, while our query FMΦ profile was derived from IPF samples, it uncovered FMΦ-like cells in many contexts, including SSc-ILD, COVID-19 lung and PDAC. These results confirm previous observations of FMΦs in lung injury^38,42^ and suggest a role for FMΦ-like cells across other organs and diseases.

### Integrated gradients analysis reveals commonalities between SCimilarity score and established gene signatures

Because FMΦ-like cells are detected by SCimilarity across many ILD studies, we hypothesized that the cells captured by different marker genes and nomenclature in different studies refer to the same biological cell state. To test this, we applied integrated gradients to quantify each gene’s importance when SCimilarity distinguishes FMΦs from randomly sampled monocytes and macrophages (**Methods**). The genes identified as important for distinguishing FMΦs are enriched in key fibrotic processes, including extracellular matrix remodeling (*MMP7*, *MMP9*, *FN1*, *SDC2*, *SPARC*, *SPP1*), lipid metabolism and lipoprotein clearance (*APOC1*, *APOE*, *LPL*, *LIPA*), and damage-associated molecular pattern recognition (*MARCO*, *MSR1*) (**Fig. 5f**, **Extended Data Fig. 4d,e, Extended Data Table 4**). While SCimilarity found many FMΦ marker genes that were already discussed in the literature, such as TREM2 (**Extended Data Fig. 4f**), it also identified novel genes elevated in FMΦs such as HLA-DQA1 and RGS1 (**Extended Data Fig. 4g,h**).

The genes with the highest importance scores in the SCimilarity embedding of FMΦs significantly overlap (p<6.7×10^-13^) with published gene signatures describing similar macrophage populations or with genes whose differential expression defined each study’s macrophage population of interest (**Extended Data Table 5**). While cell signatures from IPF lung had a high signature match (AUC ≥ 0.95), the negative control signatures of M1 and M2 macrophages^43^ had lower ones at the bottom three (AUC = 0.85 (2.65×10^-2^) and AUC=0.92 (p<4.92×10^-6^), respectively; **Fig. 5f**, **Extended Data Fig. 4d**).

### FMΦ-like cells identified among *ex vivo* stimulated peripheral blood mononuclear cells (PBMCs) help establish a novel human cell model

Research to understand the role of a novel cell state or subtype in disease, such as FMΦs, benefits greatly from the ability to model, perturb, and study the cells *in vitro.* However, there is currently no systematic way to identify *in vitro* culture conditions that generate cells that match cells identified *in vivo*. To accelerate development of an *in vitro* FMΦ system, we used SCimilarity to search for FMΦ-like cells across *in vitro* stimulated samples with the goal of identifying previously employed experimental conditions that might resemble the tissue cell state. We filtered our full reference cell collection for *in vitro* and *ex vivo* studies containing at least 50 monocytes or macrophages, resulting in 41,926 monocytes and macrophages across 40 samples from 17 such studies. These span diverse and complex conditions, such as lung organoids infected with SARS-CoV-2^44^, *ex vivo* treated acute myeloid leukemia samples^45^, or PBMCs stimulated with morphine and lipopolysaccharide^46^.

The cells most similar to our query FMΦ expression profile were monocytes grown as part of a heterogenous PBMC culture for 5 days in a 3D hydrogel culture system that was designed for expansion of hematopoietic stem cells (HSCs) from PBMCs^47^ (**Fig. 6a**, **Extended Data Table 6**). This study is unrelated to lung biology and its authors did not report any results for myeloid cells. Nevertheless, while no FMΦ-like cells were present among myeloid cells on day 0, 15% of cells grown for five or more days in this system were highly similar to FMΦs (SCimilarity score >25) and expressed *TREM2*, *GPNMB*, *CCL18* and *MMP9* (**Fig. 6b-e**). This was a surprising result, because of the seeming irrelevance of the study to fibrosis or macrophage biology and the rarity of FMΦ-like cells in PBMC samples *in vivo*.

**Fig. 6.**
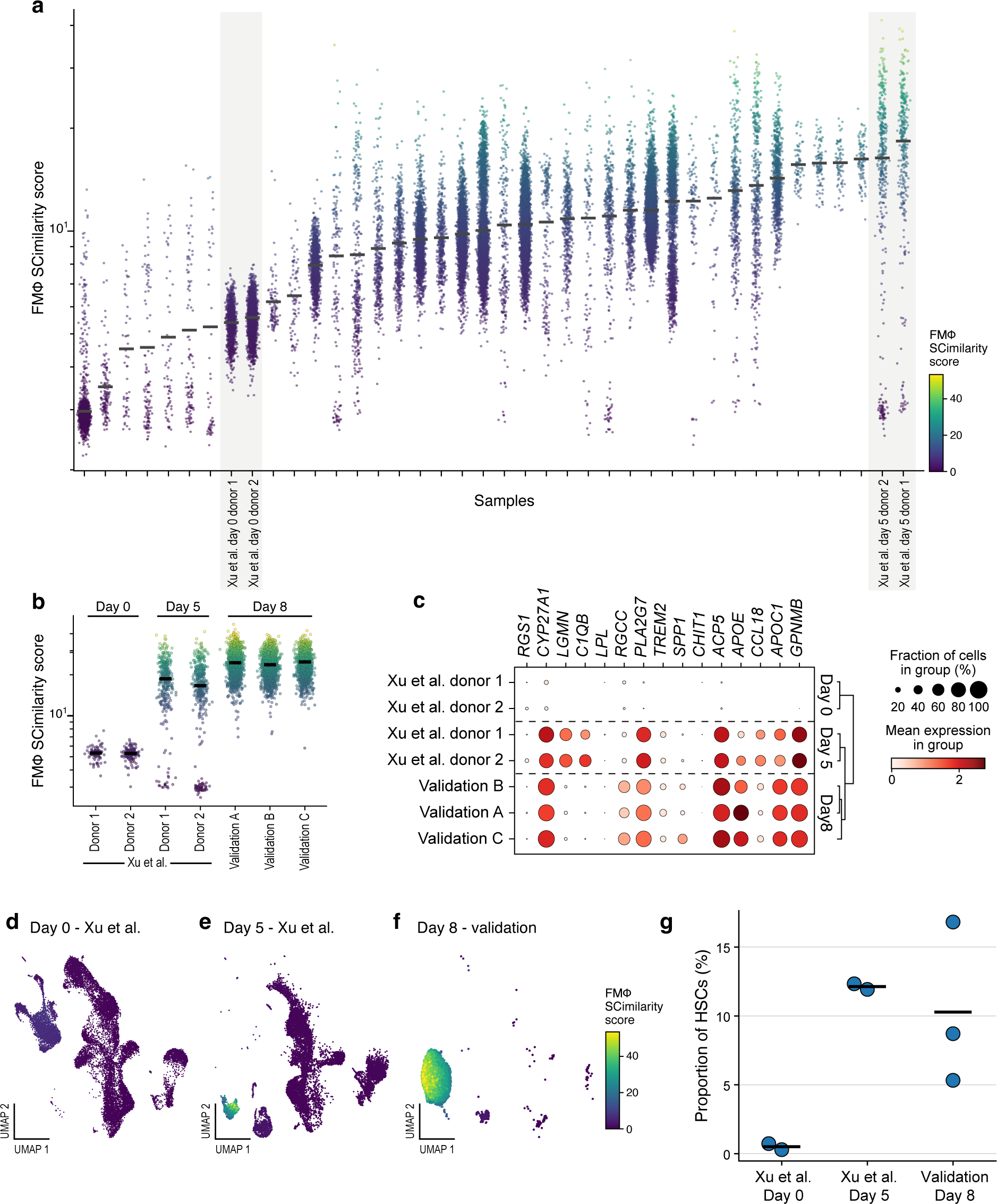
SCimilarity cell search identifies *in vitro* cells matching an *in vivo* FMΦ state and a novel *in vitro* disease model. **a,** Identification of FMΦs-like cells across *in vitro* samples by SCimilarity cell search. SCimilarity scores (y axis, log_10_ scale, and color bar) against a FMΦ query profile for each annotated myeloid cell (dot) from 40 *in vitro* samples (x axis) (from 17 studies), ordered by mean SCimilarity score. Gray boxes: Day 0 and Day 5 samples from a 3D-hydrogel culture system^4^. **b-f,** 3D conditions yield FMΦ-like cells *in vitro* in validation experiments. **b,** SCimilarity scores (y axis, log_10_ scale, and color bar) against a FMΦ query profile for each annotated myeloid cell (dot) from the original 3D-hydrogel culture system dataset^4^ and from 3 donors in the validation experiment (x axis). **c,** Mean expression (dot color) and proportion of expressing cells (dot size) of genes (rows) with high SCimilarity attribution score for distinguishing FMΦs *in vivo* (as in Fig. 5f) for myeloid cells in the original 3D-hydrogel culture system^4^ and the validation experiment (columns). **d-f,** UMAP embedding from SCimilarity’s query model latent space of cell profiles (dots) from day 0 (**d**) or day 5 (**e**) of the original 3D-hydrogel culture system^4^ or from day 8 of the replication experiment (**f**), colored by FMΦ SCimilarity score (color bar). **g,** replication of Xu et al.’s original finding of HSC expansion. Proportion of HSCs between Xu et al.’s day 0, day 5 and our validation day 8 time points.

To validate SCimilarity’s prediction of an FMΦ-like cell culture condition, we used a similar protocol to replicate the 3D hydrogel system^47^, followed by scRNA-seq to assess the yield of FMΦ-like cells (**Fig. 6b,c**,**f**). While relative cellular abundances differed between the original day 5 data (Xu et al, 2022) and our day 8 replication of the same conditions (**Methods**), 10.1% of all cells in the Day 8 experiment were predicted as HSCs by SCimilarity (**Fig. 6g**). Moreover, 41.5% of the myeloid cells in day 8 validation experiments from three donors were predicted as FMΦ-like macrophages (**Fig. 6b,f**, 37.1%, 42.5%, and 44.9%; SCimilarity score > 25). Furthermore, FMΦ hallmark genes, such as *CCL18*, *GPNMB*, *SPP1*, and *TREM2*, were enriched in the myeloid compartment of our replicate experiment compared to day 0 conditions (**Fig. 6c**). This experiment validates that an FMΦ-like population can be generated from PBMCs in culture conditions. Taken together, these results demonstrate SCimilarity’s ability to interrogate publicly available data at scale, query a reference of *in vivo* and *in vitro* data for biologically similar conditions, and help identify experimental conditions to reproduce those results in laboratory settings.

## DISCUSSION

To date, more than a hundred million human cells have been profiled across tissues in health and disease, and such data continue to grow exponentially. This growing human cell atlas should be the starting point for researchers aiming to readily search, query and compare cell states of interest across different protocols, treatments, tissues, and diseases.

SCimilarity systematically annotates and repurposes tens of millions of expression profiles from hundreds of studies, to create an integrated, searchable and queryable foundational model of pan-human cellular diversity. SCimilarity is comprised of three key features: (1) a 22.7M cell human scRNA-seq data repository (at present), (2) a foundational model for single cell gene expression with a generalizable embedding and similarity metric (which could readily be retrained for larger datasets), and (3) methods to efficiently query across this entire pan-body human cell atlas. Together, these provide new context, capabilities, and workflows for extracting insights from new and existing scRNA-seq datasets in the human cell atlas and other atlases. SCimilarity’s framework architecture can easily accommodate quick updates as data continue to grow.

Because SCimilarity can generalize to cells and datasets not seen in the training, cell profiles can be added as entirely new studies or removed by applying new cell filters without recomputing the low dimensional representations. This flexibility allows us to change the analysis’ scope at any point without redoing work, enabling modularized workflows for scRNA-seq analysis. Downstream tasks, such as cell type annotation, cell queries, and gene signature derivation all are simplified using SCimilarity’s generalized low dimensional representation and can be applied to cells not seen during training without informing the model about the importance or variability of specific genes during training. Outlier detection helps both filter out technical errors and highlight potentially novel cell subsets. Although generalized models that do not require recomputing low dimensional representations would alleviate time and expertise barriers that currently impede researchers, to the best of our knowledge, generalization has rarely been optimized in single-cell expression analysis.

There is no single objective measure of similarity, or dissimilarity, between cellular profiles. Curated gene signatures are useful when a small number of explanatory genes are sufficient to define a cell state. SCimilarity uses the full expression profile of a cell as its query, defined by either a single representative cell or the centroid of a set of cell profiles. Thus, SCimilarity’s cell-based search bypasses the manual curation requirements and biases inherent in defining a gene signature. In cases where such a gene signature is desired, SCimilarity can compute a robust signature for a cell state across studies.

Exploration of transcriptionally-similar populations across a vast atlas of human scRNA-seq data provides critical context to a cell population of interest. First, observing a query population across many similar studies shows that the original observation was reproducible, a key for subsequent scientific research^48^. Second, SCimilarity queries can connect results from independent studies. While one study may find a cell population in a disease, another may show similar cells with functional characterization, allowing us to formulate a new hypothesis on the functional properties of disease-associated cells.

This is illustrated by how SCimilarity allowed us to search for and identify FMΦ-like cells across tissues and disease states, construct a cross-study set of explanatory marker genes, and uncover a cell culture system that elicits a similar FMΦ-like state *in vitro*. Modeling FMΦs from readily available PBMCs is exceptionally valuable, because isolation of cells from human lung explants is prohibitive for many functional assays. Surprisingly, in addition to fibrotic lung, FMΦs were present in multiple tumor types, particularly PDAC, a heavily fibrotic cancer, where macrophages play an important role in mediating the associated fibrosis and have been linked to tumor progression^49^. The identification of a common FMΦ state across fibrosis, cancer, and infection suggests a broader role for these cells in the damage response and tissue remodeling processes across diseases. Moreover, SCimilarity’s search identified FMΦ-like cells in an *in vitro* study– an observation that could not have been gleaned by reviewing the paper or based on the description of the culture system – but that we validated in the lab. The variations we observed between the original and replicate *in vitro* experiment may be attributed to differences in culture duration, cell extraction from the hydrogel, lymphocyte proportions, or other batch effects. Furthermore, these results invite new hypotheses, such as whether the 3D hydrogel provides key ECM-like environmental cues that promote an FMΦ-like state and induction of remodeling genes, such as *MMP9* and *SPP1*, and which factors can be added to drive an even stronger FMΦ phenotype. Thus, SCimilarity provides a powerful framework to iteratively generate and validate such experimental hypotheses.

SCimilarity is not appropriate for all applications and will need further improvements to continue to scale with exponential data growth and to more comprehensively span human biology as the Human Cell Atlas continues to grow. Training SCimilarity requires Cell Ontology labels. Fortunately, scRNA-seq data sharing practices are increasingly relying on using the Cell Ontology for standardization. However, the Cell Ontology itself is a large, yet incomplete, effort. Cell states are only considered in training if they are recognized in the Cell Ontology, and the number of these states is growing rapidly. Furthermore, while we trained SCimilarity on vast amounts of data, cancer cells and cell lines were deliberately withheld from training due to lack of clear cell type identity and therefore may not be well represented. In addition, in our experience, we see poor performance on fetal samples, likely due to most of the training data being sourced from adult tissues.

The current data integration and cell search models provide generalizable representations of 22.7M single-cell profiles across the human body, and include a Python API for querying cell profiles of interest. Future improvements to SCimilarity could include pre-training on the massive amounts of unlabeled data, effectively exposing the model to more cell states and more technical variability during training. With effective representations we can more easily combine embeddings to include other species or data modalities. We believe that SCimilarity brings a new framework to single-cell genomics, enabling re-use of rich public data resources through instantaneous queries and demonstrates how this can be used to provide novel biological insights.

## Contributions

GH conceived of the method with input from AR, JVH, HCB, and JK. GH and TK performed data ingest and model implementation with input from JVH, ND, GS, TB, JK and AR. Python API was developed by TK with help from JVH, OS, and GH. Interpretability was developed by ND and GS with input from HCB and TB. JVH conceived of biological application of method with input from GH, ST, JR and DD. DD and TH performed experimental validation with guidance from JR and ST. GH wrote the manuscript with input from JVH, JK, AR and HCB.

## Acknowledgements

We thank Anupriya Tripathi for coming up with the name “SCimilarity” and Jenna Collier, Gokcen Eraslan, John Marioni, and Jake Freimer for their suggestions that strengthened the manuscript.

## Methods

### SCimilarity model architecture and loss function Model architecture

The SCimilarity model consists of one fully connected encoder and one decoder stage and reuses the same encoding network three times per training triplet, such that updates to the model after each batch are shared equally for each subsequent batch of training triplets. The decoder stage is not part of the conventional triplet loss architecture, but is included to compute a mean squared error (MSE) reconstruction loss.

Expression profiles are reduced through an encoder network, starting from 28,231 genes through three hidden layers with dimensions 1,024, 1,024, and 128. The 128-dimensional outputs are unit length normalized, forcing all low dimensional cell representations to lie on the surface of a hypersphere. During training, the input layer is subjected to 40% dropout, zeroing out many gene expression values at random, and each hidden layer is subjected to 50% dropout rates for maximum regularization ^1^.

While hyperspheric spaces have been infrequently used for representation of single-cell profiles ^2^, the triplet loss model often uses hypersphere embeddings to ensure consistency between the model hyperparameters ^3^. During triplet loss training, the objective is to place cells of different types sufficiently far apart. The minimum desired distance between cells of different types is called the margin. By fixing the volume of the embedding space to the surface of a unit length 64-dimensional hypersphere, the margin is interpreted consistently between model runs. Without normalization, cells can be placed up to an infinite distance apart, rendering the margin meaningless.

### Triplet loss training

To learn features that place data points considered similar near each other, the loss function depends on distances between data points embedded in a learned low dimensional latent space, described with:

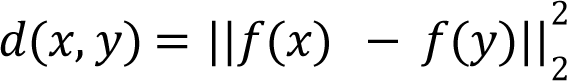

where 𝑥 and 𝑦 are two high dimensional vectors (here, cell profiles), passed through a neural network encoder 𝑓().

The triplet loss model learns from three vectors at a time: the anchor 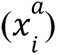, positive 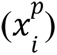 and 𝑖 negative 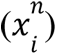. The anchor and positive vectors are considered similar, whereas the anchor and negative are dissimilar.

The model parameters are iteratively updated to decrease the number of triplets where the distance between the anchor and negative data vectors is insufficiently large relative to the distance between the anchor and the positive points, thus minimizing the triplet loss function:

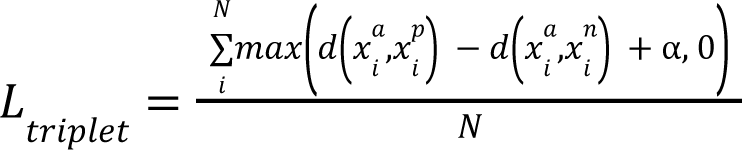

where α is the margin, which denotes how much further the negatives should be from the anchor than the positives, and *i* is the index of the triplet.

### Reconstruction loss training

The reconstruction loss is computed on the anchor cell only, because each anchor cell is used only once as an anchor within a batch. The reconstruction loss is defined as:

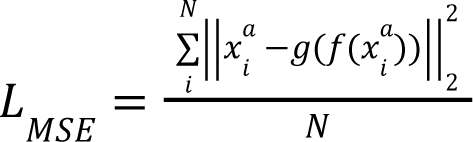

where *N* is the number of anchor cells in a batch, set to *N=*1000 in SCimilarity, and 𝑔() is the function learned by the neural network decoder stage.

### Combined loss function

Adding a reconstruction loss to classification models has been shown to improve generalization ^4^ through a regularization effect. The SCimilarity loss function combines the triplet loss and reconstruction loss functions as follows:

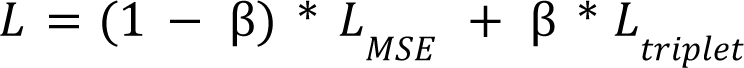

where β is a weighting term in [0, 1]. β = 0 corresponds to a conventional autoencoder, and β = 1 corresponds to a pure triplet loss model. Empirically, β = 0. 001 performed best on the cell search task (query model) and β = 1 performed best on batch integration (integration model) (Extended Data Fig. 2a).

### Use of Cell Ontology terms and relationships

Authors may annotate cell types at different granularities, which confounds triplet sampling by introducing cell type annotations with hierarchical relationships that cannot be unambiguously defined as either similar or dissimilar. As such, cell type annotations used for training are defined using standardized Cell Ontology terms and valid triplets are restricted to cells without vertical Cell Ontology relationships between members of the triplet. A vertical relationship is defined as any directed path of one or more ancestor-descendant relationships in the Cell Ontology network. Thus, there are three binary relations defined for annotation: (1) similar pairs with identical annotations (*e.g.*, “T cell” and “T cell”), (2) dissimilar pairs with non-vertical ontology relationships (*e.g.*, “CD4-positive, alpha-beta T cell” and “CD8-positive, alpha-beta T cell”), and (3) ambiguous pairs with vertical relationships (*e.g.*, “T cell” and “CD4-positive, alpha-beta T cell”). Positives are drawn from cells similar to the anchor, negatives are drawn from cells dissimilar to the anchor, and cells that are ambiguous to the anchor are excluded from sampling.

### GEO data aggregation

334 human scRNA-seq datasets were obtained from the Gene Expression Omnibus (GEO)^5^. Multiple filtering steps were used to restrict the datasets analyzed to samples from human tissue, that were generated using the 10x Chromium platform, and which reported unnormalized gene count data that could be automatically processed. To select appropriate datasets, search criteria were designed for the Biopython Entrez search tool (Cock et al., 2019) to find GEO studies that had specific properties, such as metadata keywords, file formats, and species. Then, using GEOparse^6^, the GEO text metadata was downloaded for each sample and searched for blacklisted words in the metadata or download URLs (*e.g.*, “smartseq”, “trizol”, and “fasta”) to further filter out samples that were not generated using 10x Chromium. Data for samples and corresponding download links that passed the metadata filter stage were automatically downloaded. No datasets were realigned. 753 studies were identified for download. A set of import functions was designed for the most common file type formats (.mtx, .h5ad, and gene expression matrices in .tsv or .csv). Any dataset that could not be successfully downloaded or read in was discarded. Once read in, each sample was automatically tested for count data and gene names that match a reference gene list or gene name mapper before saving each file in a uniform h5ad format for later processing. This resulted in a total of 334 published studies that were not duplicates of studies found in CELLxGENE ^7^ for use in our analysis.

### Data preprocessing

All UMI count data were natural log normalized per-cell with a scaling factor of 10,000 using the scanpy.pp.normalize_to_target(adata, 10000) and scanpy.pp.log1p(adata) functions from scanpy^8^.

### Manual data aggregation, normalization and filtering

Datasets with author-provided cell type annotations used for training were obtained from Tabula Sapiens^9^, 10x Genomics^10^, the single nucleus cross-tissue atlas^11^, and the human lung cell atlas^12^ and subjected to the same preprocessing procedures as programmatically-downloaded datasets. Cell type annotations were manually converted into terms contained within the Cell Ontology. Cells that with annotations that did not clearly map to the Cell Ontology were not included in training.

Cell profiles previously annotated as doublets, scored as doublets by infer_doublets from Pegasus^13^, had >20% total UMI counts aligned to mitochondrial genes, or had &500 total genes detected were removed.

### Preparation of training and test data

Training and test sets were chosen such that entire studies were held out of training (rather than holding out a subset of cells from each dataset) (**Extended Data Table 1**); there were 52 and 14 datasets in the training and test sets, respectively. This presents a harder generalization challenge and reflects how users are likely to use SCimilarity. Test datasets were selected to reflect the tissue diversity within the training sets.

### Selection of Cell Ontology terms for training

Cell Ontology terms were selected for training if they were observed in at least two separate studies in the training set. Terms that appeared in only one study were not used because SCimilarity is trained by comparing cells across studies. To rescue single-study terms, the immediate parent terms were inspected across studies. If a single-study term’s parent was observed in at least two other datasets then the original cell type annotation was replaced with the coarser parent term (Extended Data Table 1). Otherwise, all cells with this annotation were removed from training. As the size or annotation quality of training data grows, the number of Cell Ontology terms meeting the inclusion criteria are expected to increase.

### Triplet sampling and semi-hard triplet mining

During training, batches of 1,024 cells are sampled from the training datasets. This sampling is weighted by study and cell type to have a similar number of observations per cell type from each study per batch.

Because of the *maximum* operation within the loss function, not all viable triplets contribute to the gradient, and are categorized as easy, semi-hard or hard, based on their contribution to the gradient.

Easy negatives are defined as:

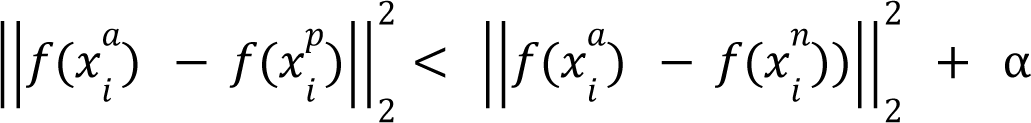

Easy negatives provide no information to the gradient because the distances between the cells in the low dimensional embedding already satisfy the objective, such that the *maximum* operation returns 0 to the triplet loss sum. Because there are many easy triplets after training a small number of batches, randomly sampling triplets does not train models effectively. To accelerate training, triplets are mined to search for training triplets that are especially informative for model training^3^.

Hard negatives are defined as:

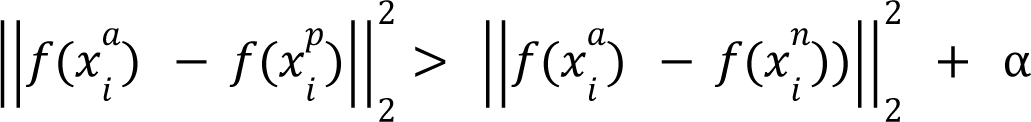

Hard negatives contribute the largest quantity to the loss function, because they do not fit and are far from fitting the desired latent relationships. In practice, hard triplets are rarely useful for training, because they contribute to model collapse during training^3,14^. Hard negatives may be enriched for incorrectly annotated cells.

Semi-hard negatives are defined as:

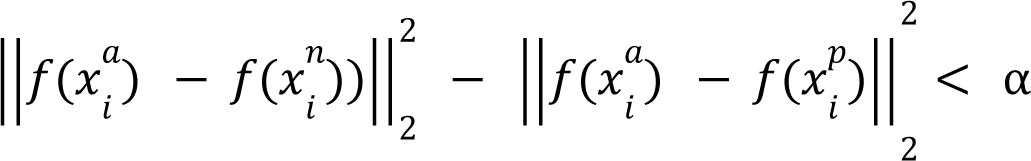

Semi-hard negatives contribute small amounts to the loss function because they nearly satisfy the desired distances between cells in low dimensional space. Meaning, the negative cell profile is further from the anchor cell than the positive cell, but by a less than desired distance α. Semihard negatives are often used in triplet loss models^3^.

Overall, we chose to train SCimilarity using only semi-hard negative triplets.

### Explainability framework and marker gene identification

An explainability framework was used to identify genes whose variation leads to the most significant variations of the learned features and, in turn, affects the relative distance between different cells.

An explanation for a pair of cells is defined as those genes which have the greatest impact on the relative distance between those cells in latent space. Given 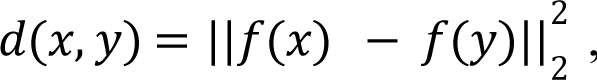, the distance between two cell profiles *x* and *y* in latent space *f*, the integrated gradient approach (Sundararajan et al. 2017) was extended to compute the importance of each gene 𝑖 in the comparison between cell profiles 𝑥 and 𝑦 as:

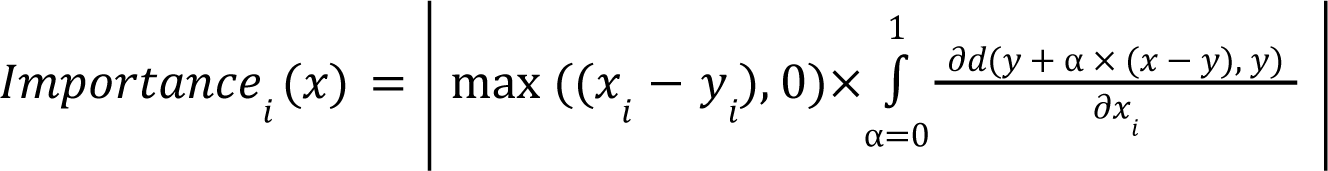

High values of 𝐼𝑚𝑝𝑜𝑟𝑡𝑎𝑛𝑐𝑒_*i*_ (𝑥) correspond to genes that are highly expressed in 𝑥, and their 𝑖 modification (i*.e.*, gradient) affects 𝑑(𝑥, 𝑦) more. Intuitively, the expression of each gene in 𝑦 is gradually increased to match 𝑥 along the trajectory from 𝑥 to 𝑦. Through this trajectory, the rate of change of 𝑑(𝑥, 𝑦) is computed for each gene, aggregating the results. The score is scaled by (𝑥_*i*_ − 𝑦_*i*_). In order to identify genes that are up regulated in a subset of interest, genes 𝑖 with expression 𝑥_*i*_ & 𝑦_*i*_ are ignored.

This approach differs in several key ways from the standard integrated gradient approach, because: (1) gradients are computed with respect to a learned distance instead of output features, (2) attributions where 𝑥_*i*_ & 𝑦_*i*_ are ignored and (3) the sign of the integral is ignored due to the complex interactions between features.

To identify important genes for a cell type 𝑡, a set of cells 𝑇∈{𝑡_1_, …, 𝑡_*N*_} with cell type 𝑡 and a set of cells with cell types different from 𝑡 are randomly sampled. Pairwise importances are computed for each pair of cells 𝑡_*i*_ in 𝑇_*j*_ and 𝑏 in 𝐵 and aggregated to obtain a signature that characterizes cell type 𝑡 as:

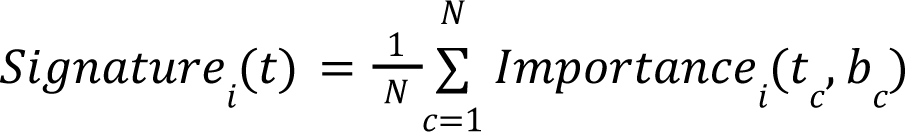

Since the pairwise comparisons are averaging relative comparisons, the sampling of {𝑏_1_, …, 𝑏_*N*_} impacts the signature scoring. To obtain general cell type markers, a background of all cell types is sampled. To obtain a cell state specific signature, a background of cells in other states of the same type are sampled. Confidence intervals for each gene 𝑖 are computed as the standard error of the mean. This results in an attribution score for each gene.

### Training and evaluation metrics

#### SCimilarity score

The SCimilarity score is defined as the inverse of the cosine distance of two embedded cell profiles:

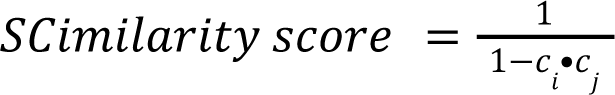

where 𝑐*_i_* and 𝑐_*j*_ are the embeddings of the *i*^th^ and *j*^th^ cell profiles with unit length, respectively and 𝑖 ≠ 𝑗. The threshold for similarity varies in practice by question and cell types.

#### Ontology-aware modified average silhouette width

Average silhouette width (ASW) has been used to assess the performance of data integration tasks on multiple scRNA-seq studies^15^ by quantifying how coherently grouped each cell type is across studies. The silhouette width of cell profile 𝑖 of cell type 𝑡 typically compares the average intra-cell type distances 𝑎(𝑖) and the average inter-cell type distances 𝑏(𝑖) between cells of type 𝑡 and cells of the nearest cell type, defined as:

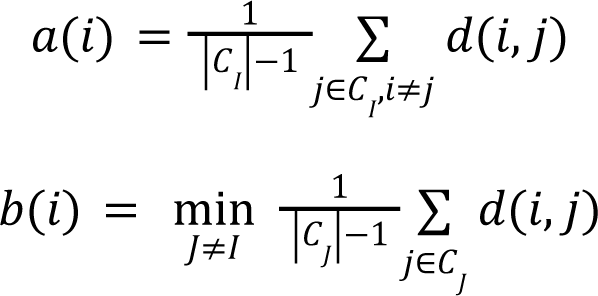

where, typically, 𝐶_*I*_ is the set of cells of author-annotated type 𝑡 and 𝐶_*J*_ are the cells of all other cell types.

However, the ASW as typically formulated does not account for differences in granularity of cell type annotations across studies. To address those, a modified formulation is used where 𝐶_*I*_ 𝐼 contains cell type label 𝑡 and all of its ontological descendants and 𝐶_*J*_ is the set of all other cell 𝐽 types, except cells of type 𝑡 and any of its ontological descendants or ancestors. For example, if computing 𝑎(𝑖) for a T cell, distances between all types of T cell terms (“CD4-positive”, “alpha-beta T cell”, “CD8-positive”, “alpha-beta T cell and CD4-positive”, “CD25-positive”, “alpha-beta regulatory T cell”, etc) are members of the “T cell” term. Ancestor terms of T cells, such as the term “Lymphocytes”, are not members of the T cell class (nor a T cell subset) but are excluded from the summation indices in the calculations of 𝑎(𝑖) and 𝑏(𝑖).

### Correlation to predefined gene signatures

To test how the SCimilarity distance represents distance between predefined cell states, a signature-based definition of cell state was correlated with the SCimilarity score (above).

For each cell in the test set, both the signature score^16^ and a SCimilarity score *vs*. the cell query are calculated, yielding two vectors, and Pearson’s correlation coefficient is calculated between the vectors.

### Selection of models for downstream analysis

Models were run in triplicate along 6 different β parameters ranging from [0,1] and one query model and one integration model were selected based on two criteria. First, query performance was tested by how well cell similarities to a query FMΦ profile correlated with a signature defining that same state (*TREM2*, *GPNMB*, *SPP1*, *CCL18*, *MMP9*, *CTSK*, *APOE*, *CHIT1*, *LIPA*, *CHI3L1*, *CD14*, *APOC1*). Second, ontology aware ASW was used to quantify how well the cells of the same type from different studies intermixed in SCimilarity’s representation. The query model was selected as the model with the highest query test performance. The integration model was selected from the β=1 models. Since the three replicates had nearly identical integration scores, we picked the model with the highest query test score as it performed much better on the query task than the other high integration models. (Extended Data Fig. 2a). The selected integration model had more study mixing than the query model according to the study (NMI) and study adjusted rand index (ARI)^15^.

### Benchmarking vs. integration methods

SCimilarity’s integration and cell search models were each compared to three batch integration methods: Harmony^17^, scVI^18^, and scArches^19^. A test dataset of 34,713 cells was created by sampling cells from lung tissue studies with uniform probability across studies. The modified ASW (above), adjusted Rand index (ARI) and normalized mutual information (NMI) were calculated as integration benchmark metrics. Harmony was run using the wrapper in Pegasus^13^ following the workflow described in https://pegasus-tutorials.readthedocs.io/en/latest/_static/tutorials/batch_correction.html. scVI and scArches were run using the scvi-tools workflow described in https://docs.scvi-tools.org/en/stable/tutorials/notebooks/harmonization.html and https://docs.scvi-tools.org/en/stable/tutorials/notebooks/scarches_scvi_tools.html, respectively. As the scArches workflow requires a reference dataset, 101,133 cell profiles were sampled across all training datasets with uniform probability across studies for use as the reference.

### Cell type annotation

Cell type assignments were performed by *k*-nearest neighbors (*k*-NN) classification combined with an annotated reference set. SCimilarity’s reduced dimensionality latent space was used to determine *k*=50 nearest neighbors in the reference data set to a query cell *t*, and the query cell was annotated either by tallying votes based each cell’s annotation with either equal weights,

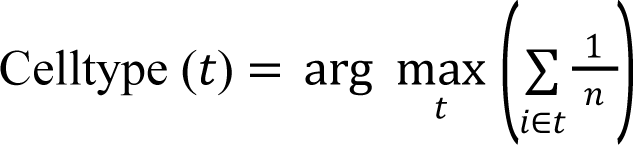

or with weights by distance in SCimilarity’s reduced dimensionality latent space:

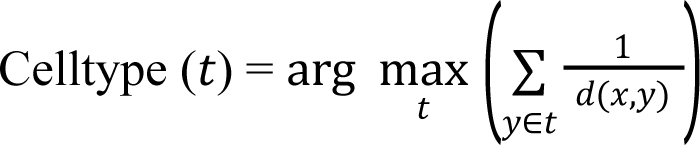

To allow users to annotate new datasets from a restricted list of cell types of interest, excluding (blocklisting) or limiting to (safelisting) specified cell type annotations is used, and is recommended when feasible to improve interpretability and reduce spurious annotations. However, extensive blocklisting or safelisting can slow the annotation process significantly, because the pre-built *k*-NN indices are not optimized for a modified target cell type list.

### *k*NN parameters for annotation and querying

Two separate *k*NN indices were used for efficient and accurate queries. For cell type annotation, a 7.9M cell *k-*NN index was built using hnswlib^20^ with ef_construction = 1000 and M = 80. Searching this *k-*NN found the 50 nearest neighbors (default behavior) for cell type annotation (k=50) and ef=100.

Cell query relied on a separate 22.7M cell *k*-NN index also built using hnswlib. This index was constructed with the following parameters: ef_construction=400 and M=50. The search parameters are set by the user’s request for how many similar cells to return. Default behavior is set to *k*=1000 and ef=*k*, but in practice *k* can vary widely depending on the use case.

### Outlier filtering

To filter outlier cells prior to visualization and downstream analysis, SCimilarity’s score is used to flag cells that are out of distribution. Cells with a SCimilarity score & 50 from the nearest cell in the training set were removed prior to further analysis. Many of these cells were from immortalized cell lines, and reflect their difference from primary cells (and absence in the training). Note that if out of distribution cells are not removed, these cells won’t be accurately annotated and can confound visualization.

### Macrophage query preprocessing

To prepare a cell query for FMΦ cells, a public dataset^21^ (GSE136831 and https://www.ipfcellatlas.com) was preprocessed with the same steps for all ingested data and scored use Scanpy’s scanpy.tl.score_genes function with a gene signature of *SPP1*, *TREM2*, *GPNMB*, *MMP9*, *CHIT1*, and *CHI3L1* Scanpy^8^. The average profile of the top 50 scoring cell was embedded using SCimilarity and used as the input query to SCimilarity’s cell search model and used throughout analyses in **Fig. 5** and **6**.

### Important genes and pathway enrichment

Important genes were identified using SCimilarity’s attribution score method. This method requires two cell groups to compare, identifying which genes differ between them. Here we used 1,000 cells that were considered similar to the average FMΦ profile calculated from Adams et al. as the FMΦ-like group. This query excluded any cells from the Adams et al. dataset. To compare to the FMΦ-like group comparison, 1,000 dissimilar monocytes and macrophages were randomly sampled (any monocyte or macrophage that was not within the top 10,000 most FMΦ similar results).

Reactome pathways enriched for the 100 genes with the top importance scores for FMΦ were determined using the method provided in the ReactomePA^22^ R package, with multiple hypothesis correction using the Benjamini-Hochberg method and the background gene universe restricted to the ∼28,000 genes included in SCimilarity. Pathways were considered significant if they met the criteria of adjusted p-value (Q) ≤ 0.05 and gene count ≥ 5.

### 3DCS culture of PBMC

Peripheral blood was sourced from healthy volunteers at Genentech that were consented as per IRB. Samples were collected in heparin collection tubes and subsequently diluted 1:1 with a solution of PBS containing 2% FBS and 1mM EDTA. 30 ml of diluted blood was overlayed onto 15 ml of Lymphoprep (STEMCELL Technologies) in a 50ml tube and centrifuged at 3,000 rpm for 20 minutes at 4°C. PBMCs were isolated from the interphase after centrifugation and diluted with PBS containing 2% FBS and 1 mM EDTA and centrifuged at 300 x g for 10 minutes at 4°C. Cell pellet was washed again with PBS containing 2% FBS and 1mM EDTA. Red blood cell lysis was performed on the cell pellet by resuspending in RBC Lysis Buffer (Cell Signaling Technology) for 5 minutes at room temperature, followed by inactivation with addition of RPMI media containing 10% FBS. Cells were pelleted by centrifugation at 300 x g for 10 minutes at 4°C and subsequently washed with PBS containing 2% FBS and 1 mM EDTA. Cells were then resuspended in a 10% sucrose solution at a concentration of 2 x 106 cells/ml right before plating into 3D hydrogel culture. Puramatrix hydrogel (Corning) was vortexed for 30 seconds and diluted 1:1 with a 20% sucrose solution. 250 µl of diluted Puramatrix hydrogel was mixed with 250 µl of resuspended PBMCs and plated in a 24-well tissue culture plate. To induce gelation, RPMI media was overlaid onto the hydrogel/PBMC mixture and incubated for 5 minutes in a 37°C incubator with 5% CO_2_. Overlayed media was aspirated off of the 3D hydrogel and washed twice with RPMI media, after which 600 µl of 3DCS media, formulated as previously described (Xu, Y. et al., Protein & Cell 2022, 13:808-824) was overlaid onto the hydrogel. Cells were cultured in a 37°C incubator with 5% CO2 for 8 days, with media exchanges every other day. On day 8, culture cells were recovered from the 3D hydrogel for scRNA-seq.

### Single cell RNA-Seq from 3DCS cultures

Wells of 3D hydrogel culture were washed with PBS, followed by recovery of the hydrogel and cells by gentle pipetting in PBS buffer. This solution was centrifuged for 5 minutes at 750 x g and the hydrogel/PBMC pellet was resuspended in TrypLE solution (ThermoFisher Scientific) and incubated at 37°C for 10 minutes. RPMI media with 10% FBS was added and the solution was centrifuged for 5 minutes at 750 x g. The resultant pellet was washed twice with PBS to remove hydrogel matrix debris. PBMCs were resuspended in PBS and passed through a 40 µM filter, pelleted by centrifugation at 300 x g for 5 minutes, and resuspended in RPMI media with 10% FBS. The cell solution was subjected to FACS to isolate cells from any remaining hydrogel debris and recovered cells were concentrated to 1,000 cells/µl in RPMI media with 10% FBS for downstream profiling by scRNA-seq.

ScRNA-seq was performed using the Chromium Single Cell 3’ Library and Gel bead kit v3 (10x Genomics), following manufacturer’s user guide. Briefly, cell density and viability of single-cell suspension were determined by Vi-CELL XR cell counter (Beckman Coulter). Cell density was used to impute the volume of single cell suspension needed in the reverse transcription (RT) master mix, aiming to achieve ∼10,000 cells per sample. cDNAs and libraries were prepared following the manufacturer’s user guide (10x Genomics). Libraries were profiled by Bioanalyzer High Sensitivity DNA kit (Agilent Technologies) and quantified using Kapa Library Quantification Kit (Kapa Biosystems). Libraries were sequenced on a NovaSeq 6000 (Illumina) following the manufacturer’s specifications with 28+90 bp paired-end reads at a depth of 101M mate-pair reads. Sequencing reads were aligned to the GENCODE 27 Basic gene model on the human genome assembly GRCh38 using Cell Ranger v6.0 (10x Genomics, Pleasanton, CA, USA).

Individual samples were genetically demultiplexing using the singularity container provided with Souporcell 2.0 ^23^. No genotype information was provided to the pipeline. Since PBMCs were provided from 3 donors, a k of 3 was used to cluster the samples into 3 genotypes. These samples were pre-processed consistently with the previously ingested samples and then embedded using SCimilarity to enable direct comparisons to Xu et al as well as the rest of the public datasets.

SCimilarity cell type classification was applied to both public and validation cells using SCimilarity with the following safelist: B cell, CD4-positive, alpha-beta T cell, CD8-positive, alpha-beta T cell, conventional dendritic cell, hematopoietic stem cell, macrophage, monocyte, natural killer cell, plasma cell, plasmacytoid dendritic cell.

### Code performance benchmarking

Benchmarks were run on servers with 8 Intel Xeon E5-2650 v4 CPUs with 2.20GHz cores and a total of 128 GB of RAM.

Query runtimes, using the pre-built approximate *k*-NN index^20^ to find the top *n* most similar cells, had an average runtime of 50 milliseconds. Some API functions use the query and summarize the metadata within one function call. That function timing is dominated by summarizing metadata and computing statistics from the query results, which requires an additional 3.3 seconds. This performance differs from an exhaustive comparison (Fig. 5b), where the query was directly compared against 2.58M monocytes and macrophages with a runtime of 2 seconds.

Cell signatures were calculated using scanpy.tl.score_genes. The scanpy score_genes function was applied to the already normalized data. This runtime totalled 2 hours, 46 minutes and 20 seconds when it was applied across each h5ad file (one file per tissue sample). Even though h5ad files were not stored with any compression, file reading was a dominant factor in runtime.

### Code availability

Code and tutorials are available at https://github.com/Genentech/scimilarity.

### Licensing

- Code license: Apache 2.0
- Pretrained model weights, kNN and pre-built indices license: CC-BY-SA 4.0

## Extended Data figure legends

**Extended Data Fig. 1.**
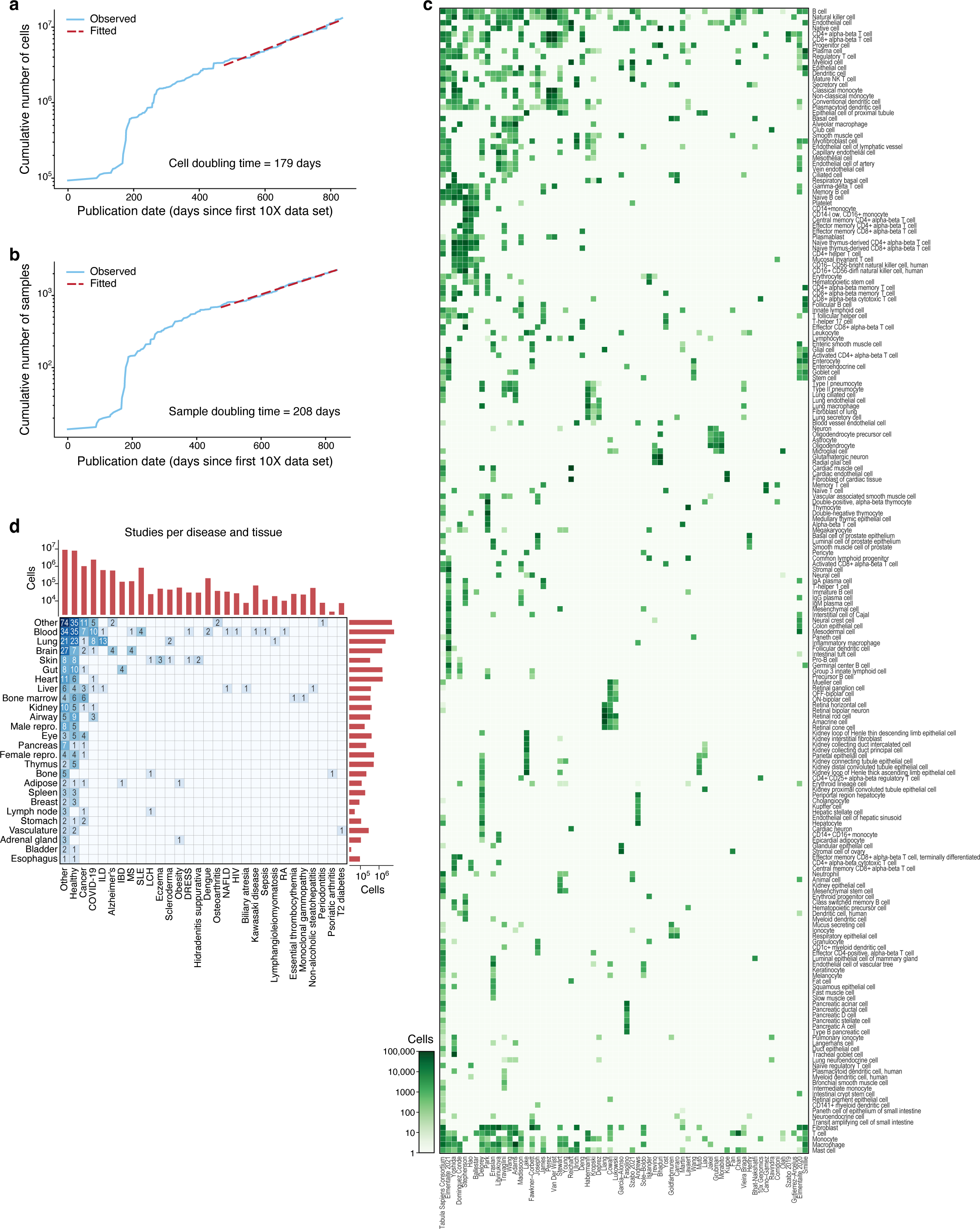
Data compendium to assemble a pan-human reference. **a,b**, Cumulative number of cells (**a**, y axis) and samples (**b**, y axis) profiled by sc/snRNA-seq (and matching our filters; **Methods**) over time (x axis). Doubling time is calculated based on the publication date from the most recent 150 data points (dashed red line). **c,** Author-annotated cell types used in training. Number of author-annotated cells (color bar) from each Cell Ontology type (rows) and study (columns) used for SCimilarity model training. **d,** Tissues and diseases used in training. Number of studies (heatmap tiles, text and color bar) and cells (margins, y or x axis) used for model training from each tissue (rows, right y-axis) and disease (columns, top x-axis).

**Extended Data Fig. 2.**
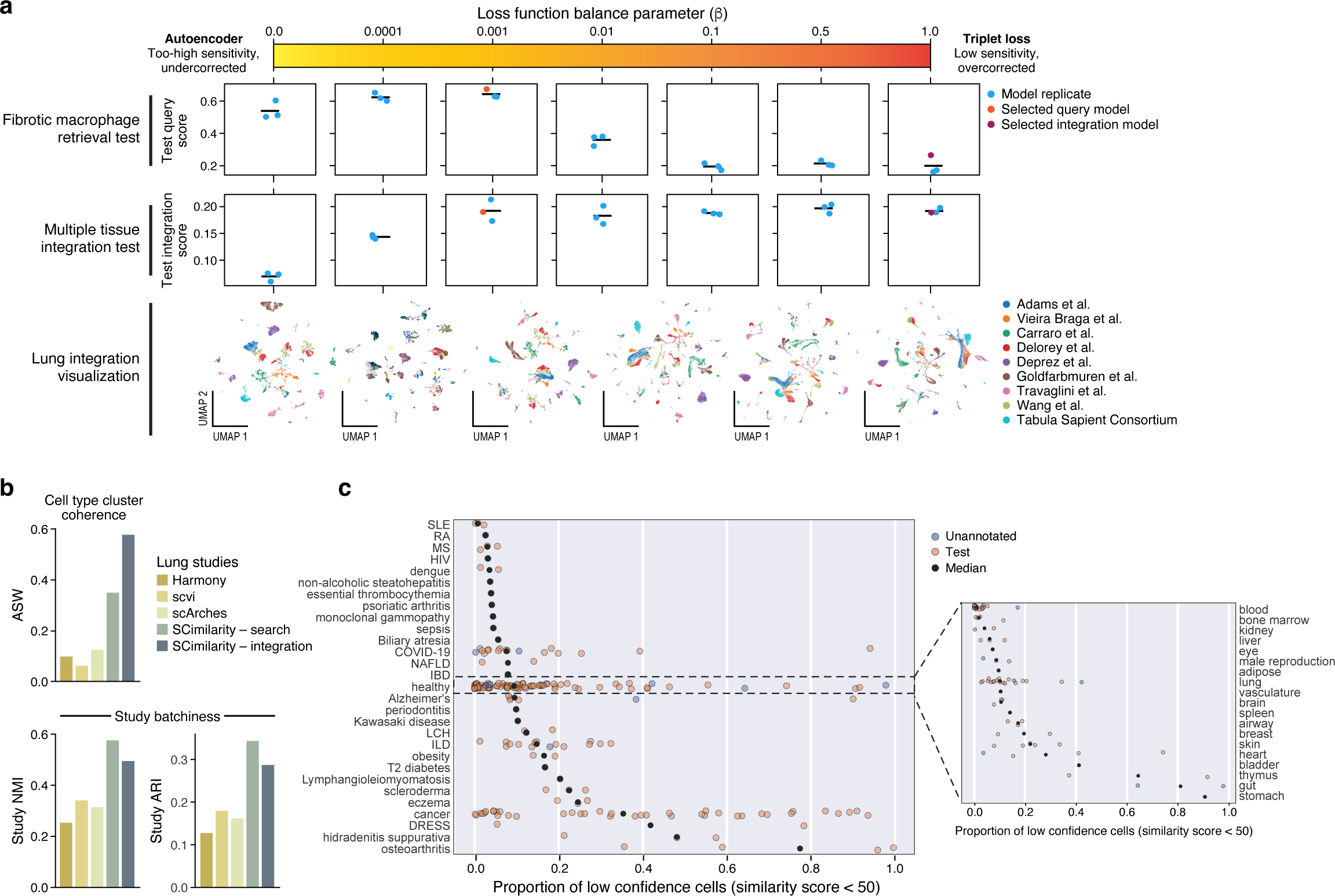
SCimilarity training details and hyperparameter search. **a,** Impact of triplet and autoencoder loss mixing on model performance, where the leftmost column is a traditional autoencoder and the rightmost column is exclusively triplet loss. The FMΦ retrieval test quantifies how much correlation there is between signature scoring of FMΦs and SCimilarity score to FMΦ. The multiple tissue integration test quantifies an ontology-aware average silhouette width where a higher score denotes more coherent clusters for each cell type. The bottom row shows UMAPs for each loss function mix for nine lung scRNA-seq datasets, colored by study. **b,** Benchmarking SCimilarity to established data integration models. Ontology-aware average silhouette width (ASW, y axis, top), normalized mutual information (NMI, y axis, bottom left) and adjusted Rand index (ARI, y axis, bottom right) for SCimilarity’s integration and search models and for scVI, scArches, and Harmony (x axis), each applied to nine lung datasets. **c,** Outlier cells from different types and tissues. Fraction (x axis) of cells from different disease (left) or healthy (right) tissue samples with low similarity (SCimilarity score <50) to training data.

**Extended Data Fig. 3.**
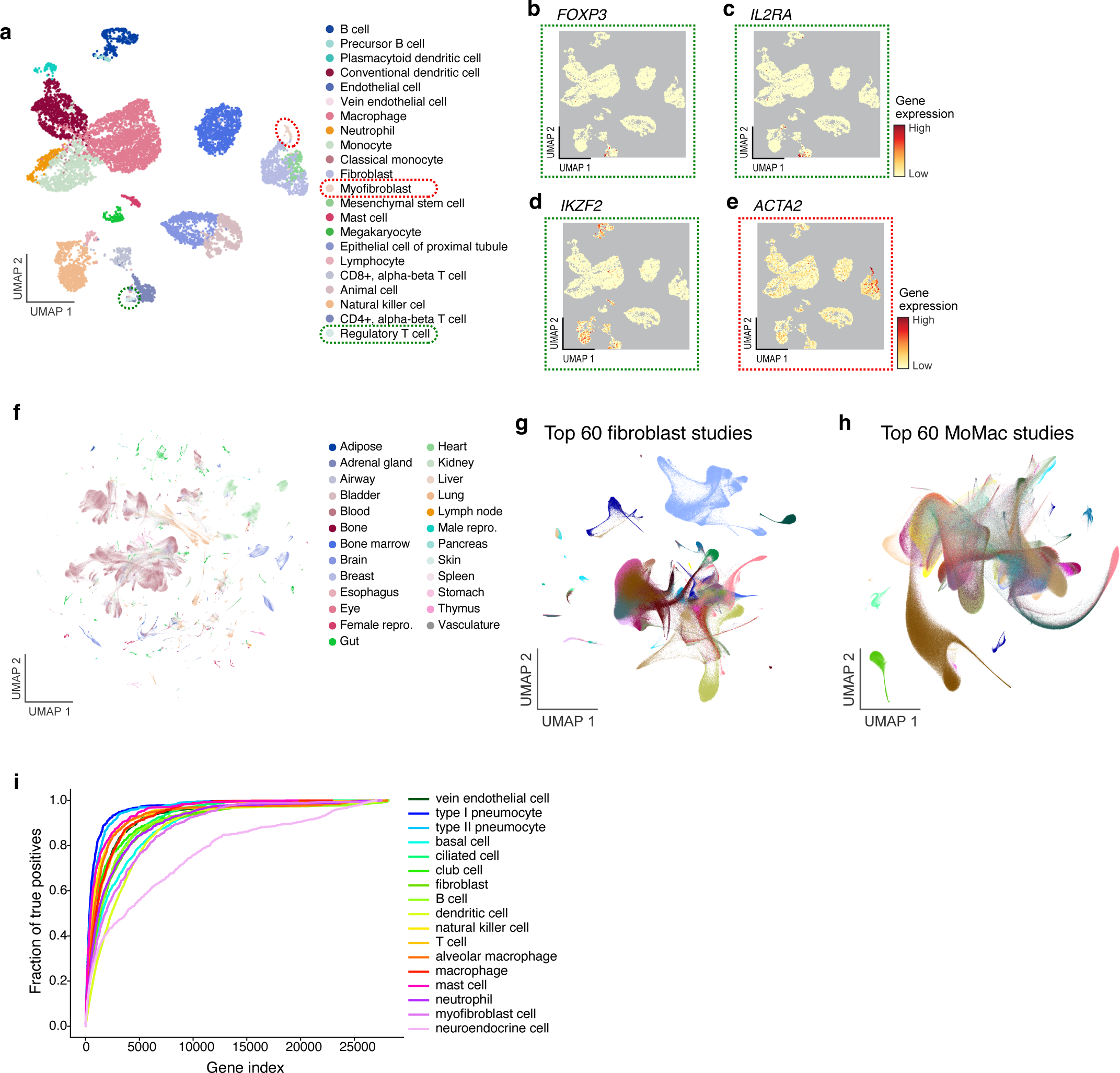
Validation of large-scale integration and annotation. **a,** Unconstrained cell annotation. UMAP embedding of single cell profiles (dots) from SCimilarity’s latent representation of the held out scRNA-Seq kidney data^1^ (as in Fig. 3b**,c**, colored by cell annotation without constraining target labels to the scope of author-provided labels in this studyor by expression of select marker genes of regulatory T cells (**b-d)** or myofibroblasts (**e**). **f,** UMAP of 2,000,000 embedded cells uniformly sampled from the 22.7M reference, colored by tissue (as in Fig. 4b). **g,h,** UMAP embedding of cell profiles predicted by SCimilarity as fibroblast/myofibroblast (**g**) or monocytes/macrophages (**h**), colored by study (for the 60 studies contributing most cells). **i,** SCimilarity cell-type important genes match cell-type specific signatures. Fraction of cell type-specific differentially expressed genes (from Eraslan *et al.*^5^) (y axis) captured by top-n important genes (x axis) for that cell type by SCimilarity’s integrated gradients attribution analysis.

**Extended Data Fig. 4.**
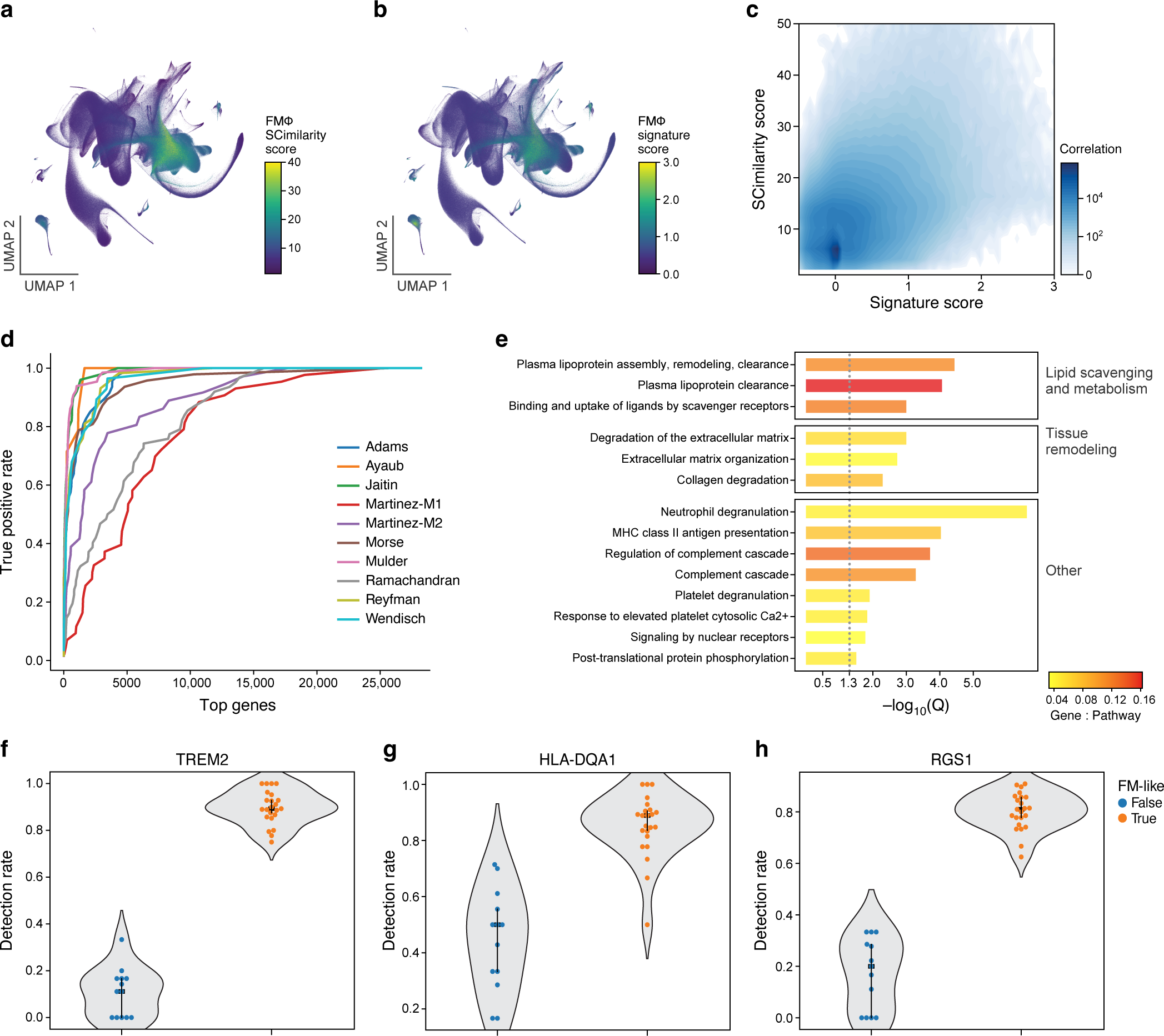
FMΦs among monocytes and macrophages. **a-c,** Agreement between SCimilarity and traditional FMΦ cell scores. **a,b** UMAP embedding of 2,578,221 monocyte and macrophage cell profiles (dots) from SCimilarity’s latent space representation colored by SCimilarity score using a prototypical FMΦ cellular profile defined from Adams *at el*.^6^ (**a**) or Scanpy’s signature score for FMΦ associated genes (**b**). (**c**) Scanpy FMΦ gene signature score (x axis) and FMΦ SCimilarity score (y axis) for each cell (shown as density). **d,** Agreement between SCimilarity FMΦ important genes and published FMΦ signatures. ROC curve of the fraction of each study’s gene sets (y axis) captured within the top genes by SCimilarity attribution ranking (x axis). **e,** FMΦ important genes are enriched for relevant pathways. False discovery rate (-log_10_(q value), hypergeometric test, x axis) for enrichment of Reactome pathways (y axis, Q ≤ 0.05 and gene count ≥ 5) with the 100 genes with the top integrated gradients attribution scores for the FMΦ query (ranked by score). Color: ratio of important genes within a Reactome pathway to the total size of the pathway. **f-h,** Expression of known and novel genes associated with FMΦs. Pseudobulked gene expression values for ILD tissue samples for known marker TREM2 (**f**) and enriched genes not previously described (**g,h**).

